# DeepTMHMM2 enables accurate prediction of transmembrane protein topology and subcellular location

**DOI:** 10.64898/2026.08.24.746435

**Authors:** Felix Teufel, Jeppe Hallgren, Henrik Nielsen, Anders Krogh, Konstantinos D. Tsirigos, Ole Winther

## Abstract

Transmembrane α-helical and β-barrel proteins are a ubiquitous component of proteomes. Topology prediction infers how proteins are embedded in lipid bilayers, identifying membrane-spanning segments and their orientation. While recent methods achieve high performance for membrane-spanning segments, they cannot predict re-entrant regions and interfacial helices – membrane-associated segments that partially insert but do not cross the bilayer – nor identify which biological membrane a protein resides in. Here, we present DeepTMHMM2, the first predictor to include re-entrant regions and interfacial helices in its topologies and jointly predict localization across 17 biological membranes. Benchmark results show that DeepTMHMM2 successfully learns to predict the additional elements, while achieving strong performance on canonical α-helical and β-barrel topology prediction. Applying DeepTMHMM2 to Swiss-Prot reveals that non-crossing segments are a ubiquitous feature of the transmembrane proteome, with interfacial helices present in nearly a quarter of all α-helical transmembrane proteins.

## Main

Approximately one third of proteins in a proteome are transmembrane (TM) proteins, which play central roles in a wide range of cellular processes^1^. Important classes of TM proteins, including G-protein coupled receptors (GPCRs), transporters, and ion channels, are therefore of considerable interest as drug targets^2^. Despite their importance, TM proteins have remained underrepresented in the Protein Data Bank (PDB)^3^ since their structural characterization is difficult due to challenges associated with crystallization.

Over recent years, Cryo-EM has made experimental structure determination more tractable^4^, and highly accurate prediction methods such as AlphaFold^5,6^ have shown strong performance on TM proteins^7^, substantially increasing the availability of structures. However, while a 3D structure captures the relative positioning of TM segments in space, it does not directly report on the orientation of the protein as a whole with respect to the membrane. By providing this orthogonal information on the individual residue level, sequence-based transmembrane topology prediction methods have maintained their relevance in the age of structure prediction, with combined predicted structure-topology workflows now supporting applications in de novo antibody design^8^, virology^9^ and target discovery^10,11^.

Topology prediction can be traced back to the early 1990s, with TopPred being the first method to tackle this task^12,13^. TMHMM was one of the earliest and most widely used methods since its launch more than 25 years ago^14,15^, though it was limited to α-helical TM proteins (TM-α), excluding signal peptides (SPs) and β-barrels (TM-β). Numerous other methods have followed, making use of various machine learning approaches, like hidden Markov models (HMMs), neural networks (NNs), support vector machines (SVMs) and dynamic Bayesian networks (DBNs)^16^. Some of the methods used only the protein sequence as input, others a multiple sequence alignment (MSA), and others combined multiple predictors into one consensus output^17–19^. With the advancement of protein language models and deep learning, embedding-based methods like DeepTMHMM^20^ and TMbed^21^ have become the state of the art.

While these advances have led to high performance for α-helical and β-barrel TM segments as well as SPs, less prevalent membrane-associated elements – including re-entrant regions (RE), which penetrate but do not fully traverse the membrane, and interfacial helices (IF), which associate laterally with the membrane surface – remain difficult to predict. A few methods such as OCTOPUS^22^ and MEMSAT-SVM^23^ included RE prediction, achieving only limited performance on very scarce annotated data, whereas MemBrain 3.1^24^ additionally included IF by relying on computationally inferred labels for training.

Here we introduce DeepTMHMM2, which extends topology prediction to all major membrane-associated segment types in TM proteins (Figure 1a). In addition, DeepTMHMM2 classifies the biological membrane type, discriminating among 17 membranes (Figure 1b). While labeling sides as the inside and outside of a generic membrane, as done in previous topology predictors, is easily interpretable for cell membrane proteins, these terms become ambiguous when including organelles. As an example, the inside of the ER, the lumen, is topologically equivalent to the outside of the cell when tracing a protein through the secretory pathway. By predicting membrane types, DeepTMHMM2 always assigns interpretable names to the two sides.

**Figure 1.**
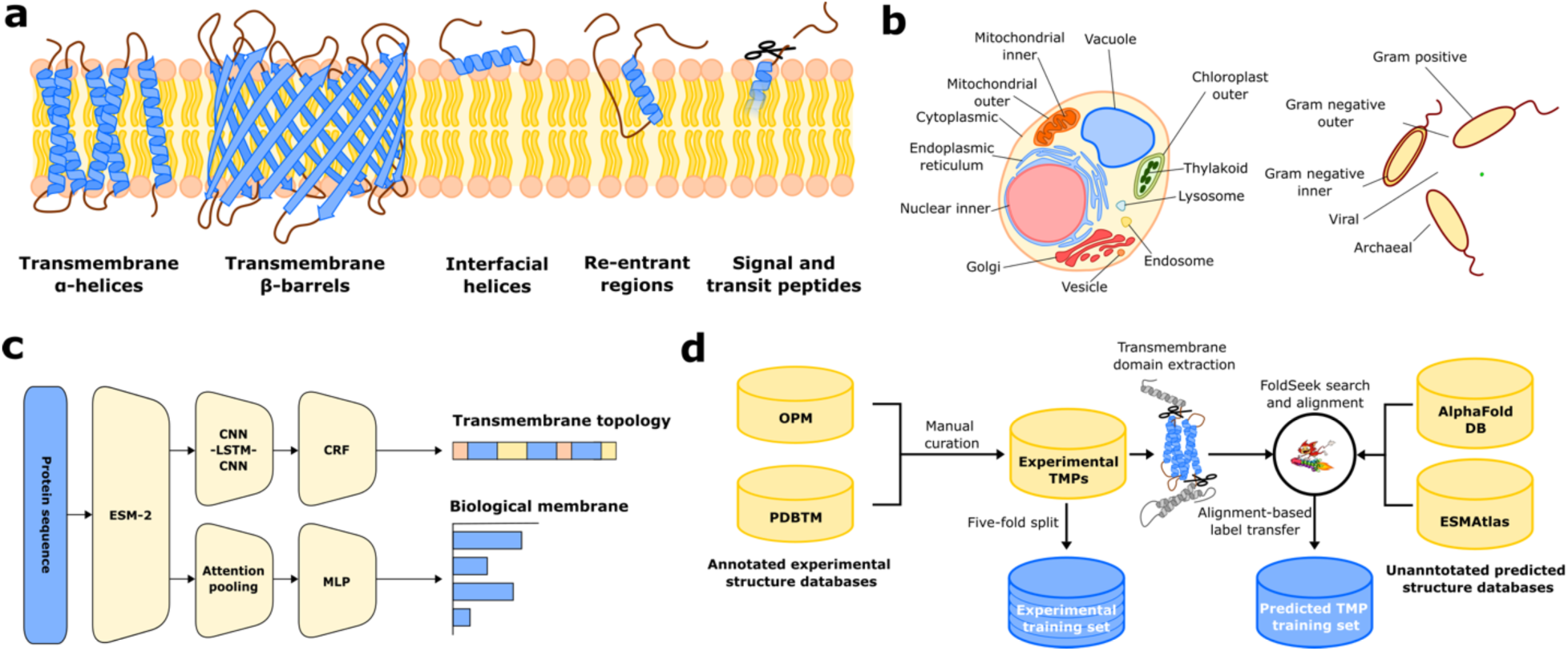
Modeling transmembrane protein topology. **a.** The six transmembrane protein topology features predicted by DeepTMHMM2. **b.** Overview of the 17 biological membrane types predicted by the model. **c.** Model architecture. DeepTMHMM2 combines two models, a CRF for protein topology and a multiabel classifier for membrane types. **d.** Topology training data generation strategy. Starting from a manually reviewed dataset of annotated experimental protein structures, FoldSeek is used to identify and annotate predicted structural homologs.

DeepTMHMM2 is a sequence-to-sequence model. It embeds amino acid sequences using the ESM-2 protein language model^25^ and extracts features using a Convolutional - Long Short-Term Memory Neural Network (CNN-LSTM). A conditional random field (CRF)^26^ decodes residue-level predictions while enforcing biologically valid topologies (Figure 1c). Moreover, as the CRF is a probabilistic model, it enables learning from partially observed experimental data (Methods). Membrane types are predicted from the ESM-2 embeddings by a separate attention-based classifier head^27^ in parallel to the topology. As a single protein can localize to multiple membranes, the classifier performs multilabel prediction.

We compiled a dataset of 14,812 soluble, 3,191 TM-α and 221 TM-β proteins, including SPs, transit peptides (TPs), TM segments, RE and IF in the topologies. Transmembrane annotations were obtained from the experimental databases OPM^28^ and PDBTM^29^, and combined with additional information from UniProt^30^ (Methods). To establish sufficient data for training, we manually curated RE and IF from literature and structural evidence, using PDBTM annotations as the starting point (Supplementary Note 1). In total, 540 and 98 proteins were annotated with IF and RE elements after curation.

We reduced the data to 20% maximum pairwise sequence identity, leaving 8,094 sequences (7,222 soluble, 780 TM-α and 92 TM-β, Tables S1-S2) and partitioned the dataset into five folds. To further enrich the training data, we developed a pipeline that uses Foldseek^31^ to search the AlphaFold Protein Structure Database^32^ and the ESM Metagenomic Atlas^25^ for highly sequence-divergent structural homologs of the experimental TM-β and TM-α proteins, transferring topology annotations to remote homologs using structural alignment (Figure 1d). This yielded 9,660 TM-α and 5,316 TM-β proteins with maximum 20% identity to any sequence in the five folds. To aid membrane type prediction, we prepared an additional dataset of 13,277 membrane proteins from UniProt with known subcellular location but undetermined topology at 20% identity (Table S3).

We trained and evaluated DeepTMHMM2 in five-fold cross-validation, using the structural homolog dataset as auxiliary data that was mixed into the topology training batches. We evaluated performance for each protein class, reporting both classification and topology prediction performance. A topology counts as correct when all TM segments, orientations and any sorting signal are predicted correctly. For SPs and TPs, this criterion corresponds to predicting the cleavage site (CS). RE and IF are evaluated separately from the topology correctness criterion to allow comparison with existing methods. We trained the membrane type classifier separately in cross-validation, mixing in the additional UniProt dataset during training.

For benchmarking, we use available cross-validated predictions of DeepTMHMM1, TMbed and SignalP 6.0 to construct common validation sets with DeepTMHMM2. We can thus compare performance without the need for retraining. We also benchmark prediction methods that do not have cross-validated data available in Tables S10-S13.

## Results

### Topology and membrane type prediction performance

For classification, DeepTMHMM2 reaches a performance level that is similar to DeepTMHMM1 and TMbed for α, β and SP, with differences often lying within or close to the margin of error (Figure 2a, Tables S6-9). TMbed retains an edge in F1 score on α (0.960 vs 0.977) and SP (0.962 vs. 0.985), driven by a reduced number of false positive α predictions (precision 0.942 vs 0.997), whereas DeepTMHMM2 operates at a trade-off more biased towards sensitivity for α (recall 0.979 vs 0.968), and precision for SP (precision 0.997 vs 0.985, recall 0.930 vs. 0.985). We conclude that core classification (α, β and SP) performance may be near saturation for these three methods of similar architecture, subject to method-specific multi-objective trade-offs between classification and topology across types.

**Figure 2.**
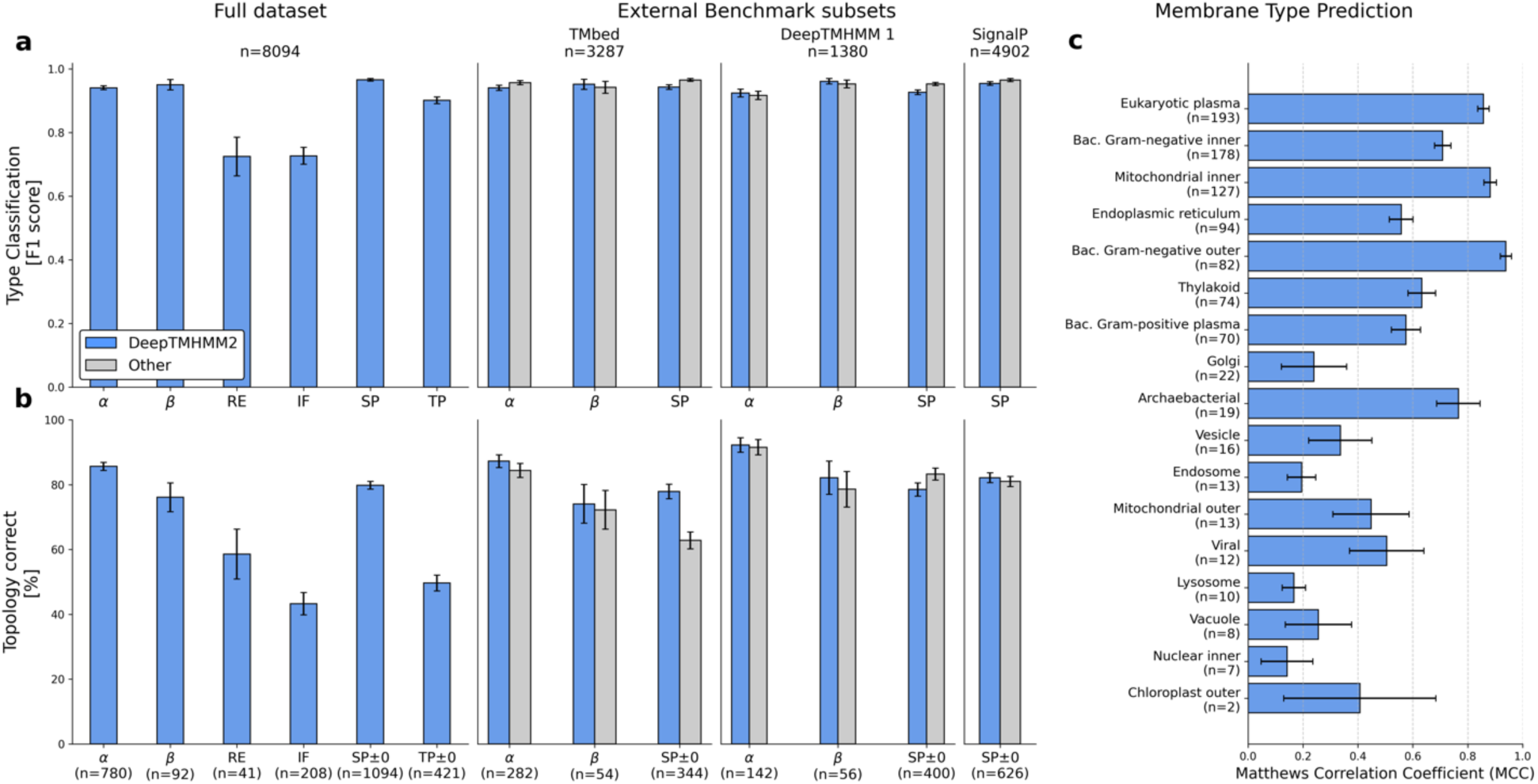
Prediction performance of DeepTMHMM2. We evaluate performance on the full cross-validated dataset, and additionally on three two-way homology reduced subsets to allow benchmarking against TMbed, DeepTMHMM1 and SignalP 6.0. **a.** Protein type classification performance. **b.** Topology prediction performance. **c.** Membrane type prediction performance.

For detection of proteins containing RE and IF, DeepTMHMM2 exhibits F1 scores above 0.7, showing that the model succeeds at predicting the two additional features. The F1 scores are dominated by high precision (0.89 and 0.84), with the recall remaining modest (0.61 and 0.64). The baselines (MEMSAT-SVM for RE and MemBrain 3.1 for IF) reach F1 scores of 0.38 and 0.31 (Tables S13-S14). For TP detection, the F1 score of 0.90 is similar to the reported performance of TargetP 2.0^33^.

DeepTMHMM2 reaches 85.6% and 76.1% correct topologies for TM-α and TM-β (Figure 2b). Despite supporting a broader range of topological features, its performance on the benchmark subsets remains comparable to that of TMbed and DeepTMHMM1 for both TM types, with DeepTMHMM1 performing particularly strongly on SP prediction. We observe that TMbed underperforms at predicting the exact CS position, but recovers performance with increasing tolerance (Figure S2). The performance of SignalP 6.0 and DeepTMHMM2 is comparable. For TPs, performance at tolerance 0 stays modest, but improves starting from tolerance 1 (Figure S3). This is expected, as e.g. TargetP uses a five-residue tolerance on CS positions to account for annotation noise.

Even though the training data is limited, DeepTMHMM2 learns to predict the positions of RE and IF at 58.5% and 43.3% correct. The performance on this metric is a consequence of the classification recall, which indicates that only 61% of RE-containing and 64% of IF-containing proteins have an element predicted at all, putting an upper bound on topology correctness. For these proteins that do receive an RE or IF prediction, the positioning is correct in 96% and 68% of cases, respectively.

We find strong membrane type prediction performance for the cell membranes of eukaryotes, bacteria and archaea (Figure 2C, Table S16). For organelles, performance is more varied, with good performance for the mitochondrial inner, endoplasmic reticulum (ER) and thylakoid membranes. Several other organelle membranes remain hard to predict, including the Golgi (MCC 0.24), Vacuole (0.26), Lysosome (0.17), and Nuclear inner (0.14), each with few positive examples in the cross-validation dataset.

### DeepTMHMM2 predictions reflect membrane biology

Biological membranes differ in their lipid composition, causing differences in their thickness. Given cross-validated topology predictions grouped by their true membrane type, we investigate whether prediction behaviour is in agreement with known properties (Figure 3a). While topologies do not directly predict thickness, we can use TM-α segment length as a proxy, which is commonly understood to be correlated with thickness^34^.

**Figure 3.**
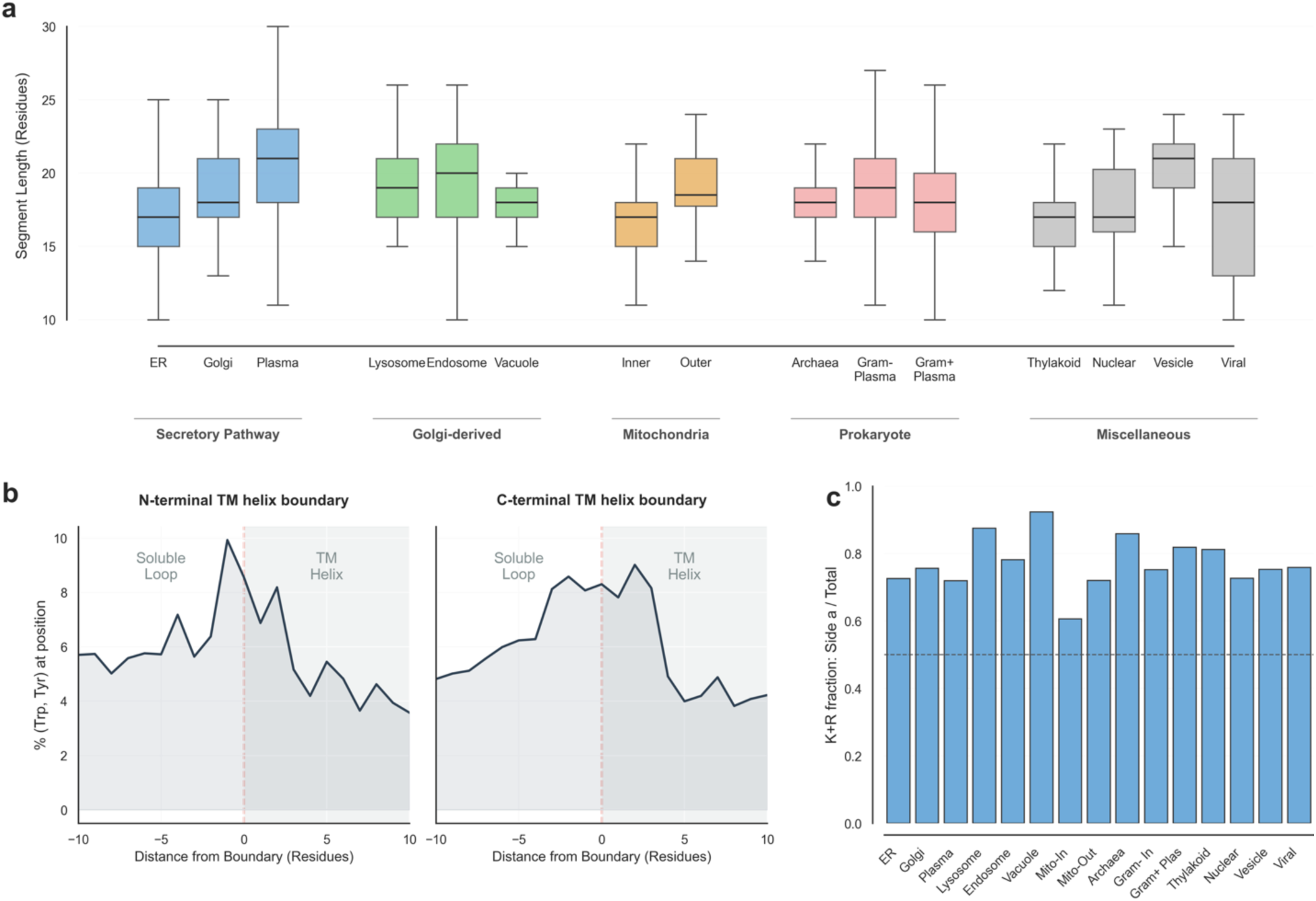
Predictions capture the biophysics of membrane insertion. **a.** Distribution of predicted TM-α segment lengths by membrane type. **b.** Frequency of aromatic residues (Trp, Tyr) relative to the N- and C-terminal boundaries of predicted TM segments. **c.** Fraction of positively charged (Lys, Arg) residues in TM-flanking segments on the topological side *a* (inside).

Eukaryotic cells exhibit a thickness gradient from the ER to the cell membrane, driven by an increasing cholesterol concentration^35,36^. Model predictions recapitulate a gradual increase of average segment length from the ER to the cell membrane via the Golgi (One-sided Mann-Whitney-U tests, p<0.001). We also find that the trans-Golgi derived lysosome, endosome and vacuole membranes have a similar segment length to the Golgi. The cardiolipin-rich mitochondrial inner membrane is understood to be one of the thinnest membranes^37^. In our predictions, we find it to be slightly thinner than the ER membrane (p<0.05). However, the topology predictions fail to reproduce that the mitochondrial outer membrane is thinner than the inner membrane. This specific comparison may be unreliable as only 8 TM-α proteins are annotated to localize to the outer membrane (inner: 120), and, 3 out of these 8 localize to multiple membranes.

Beyond matching expected thickness properties, DeepTMHMM2 correctly resolves the physical principles of the lipid-water interface. While the exact boundary placement in structure-derived annotations is subject to uncertainty, it is known that Tryptophan and Tyrosine have a natural tendency to occur at the membrane edge to anchor the protein, forming what is termed an “aromatic belt” ^38–40^. Consistent with this expectation, we find Trp and Tyr enrichment on both the N-terminal and C-terminal predicted helix boundaries (Figure 3b).

Finally, we investigate the well-known positive-inside rule, which states that Arginine and Lysine are more prevalent in cytosolic than in non-cytosolic loops. This rule was first observed in the bacterial inner membrane^41^, and later in eukaryotes and archaea^42,43^, serving as a direct component of earlier topology prediction methods^13,44^. We count the number of Arg+Lys in predicted loops on both sides, reporting the count on side *a*, divided by the total count (Methods). We consistently find mean ratios >0.5, indicating positive-inside enrichment across all membranes (Figure 3c). Interestingly, this bias is most pronounced for lysosome and vacuole membranes, where the typically acidic luminal pH may impose a stronger selection against positively charged residues.

### Understanding the biological role of re-entrant regions and interfacial helices

We next applied DeepTMHMM2 to 574,627 proteins in Swiss-Prot, yielding 81,732 TM-α and 1,775 TM-β predictions (Figure 4a). Given the high precision achieved for RE and IF, we investigated the occurrence of these elements. We find RE and IF to be remarkably prevalent, occurring in 3.8% and 24.9% of all TM-α proteins. The incidence in TM-α proteins in the human reference proteome is similar, at 5.3% and 23.1%. To gain further insight into the biological role of proteins with RE and IF, we performed Gene Ontology (GO) enrichment analysis, comparing proteins with RE/IF features against the background of all TM-α proteins, reporting gene ratios (GR, the fraction of feature-positive proteins having the given GO term) and enrichment (fold change over the TM-α background prevalence). Findings reported in the following all have an adjusted p-value<0.01. Across all of Swiss-Prot, we find that RE are mainly enriched in transport and pore-forming mechanisms (Figure 4b). Monoatomic ion transmembrane transporters account for over half of all proteins predicted to contain RE (enrichment = 2.15, GR = 0.58). Furthermore, RE are strongly associated with gated ion channel activity (enrichment = 6.95, GR = 0.24), where RE can form elements of the pore such as the selectivity filter^45^.

**Figure 4.**
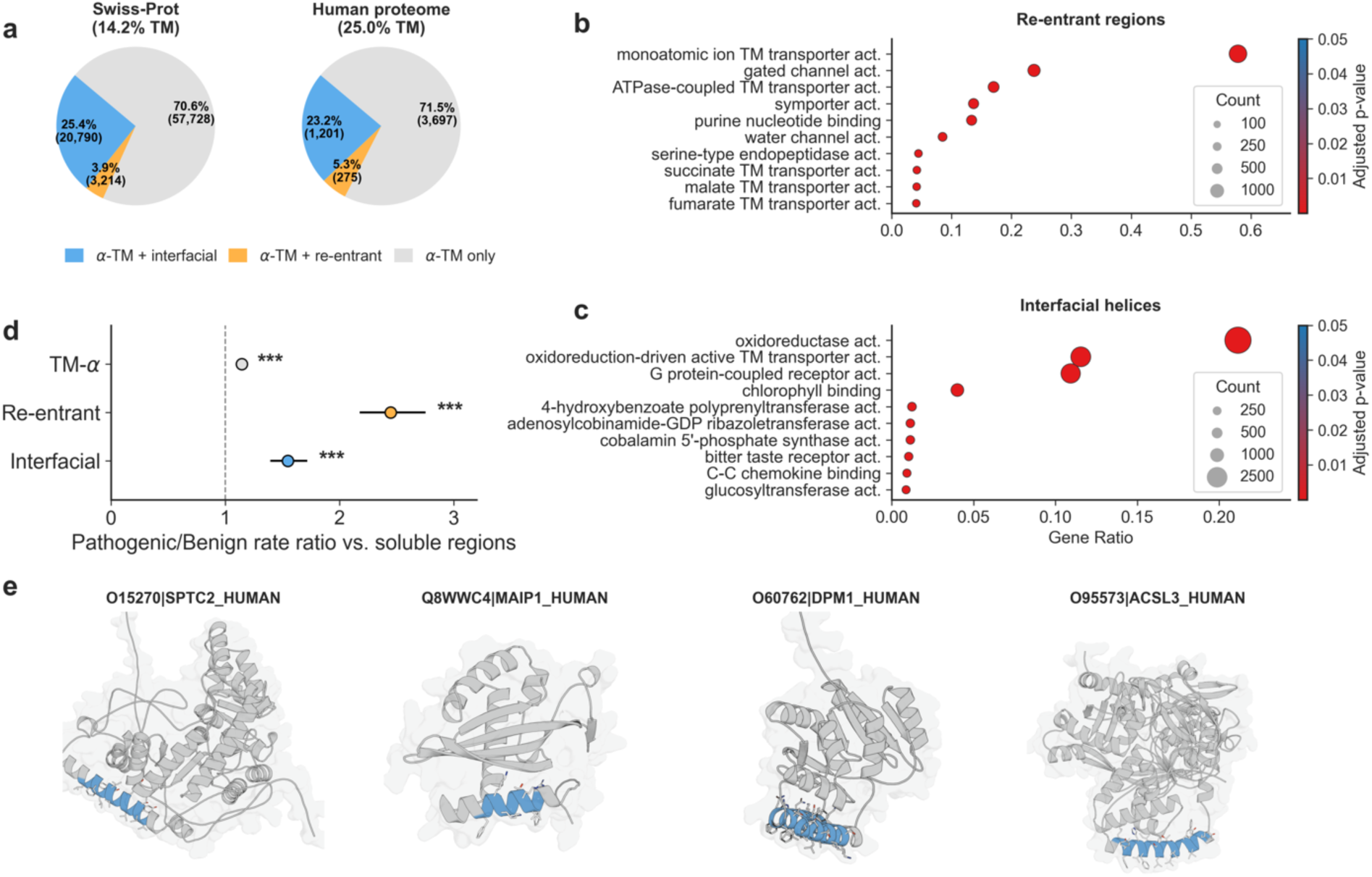
RE and IF segments are ubiquitous and functionally important. **a.** Membrane protein types predicted by DeepTMHMM2 in Swiss-Prot and the human reference proteome (Release 2026/01). **b and c.** Gene ontology (GO) term enrichment analysis for RE and IF-containing proteins in Swiss-Prot. The 10 terms with lowest adjusted Fisher’s exact test p-value are shown, sorted by GR. **d.** Enrichment of pathogenic ClinVar variants in TM-α, RE and IF segments of human TM-α proteins vs. soluble segments. **e.** AlphaFold predictions of peripheral membrane proteins with DeepTMHMM2 predicted interfacial helices highlighted.

IFs are enriched in proteins involved in signaling and energy transduction. Specifically, we find IF in GPCRs (enrichment = 2.08, GR = 0.11), where they correspond to the C-terminal amphipathic helix present in some types of GPCRs^46^ (Figure 4c). Energy-transducing systems with IF enrichment include oxidoreduction-driven active TM transporters (enrichment = 3.65, GR = 0.12) and chlorophyll-binding proteins (enrichment = 2.09, GR = 0.04). More broadly, 41% of the annotated oxidoreductase proteome in Swiss-Prot is predicted to possess at least one IF (enrichment = 1.27, GR = 0.21). Finally, we also observe significant enrichment of several distinct transferase activities.

We further investigated the functional importance of RE and IF in the human proteome by analyzing pathogenic variants in ClinVar^47^ (Figure 4d). After correcting for segment length, pathogenic variants are significantly enriched in RE, IF and TM-α compared to soluble regions in TM proteins (Poisson GLM, p<0.001, Methods). The rate ratios for both RE and IF exceed that of TM-α, highlighting that such elements can perform function-critical roles. However, we note that as the availability of ClinVar annotations is highly biased towards a small number of well-studied proteins^48^, this result is to a large degree driven by mutations in ion channel proteins.

### Interfacial helices in TM proteins enable transfer learning to peripheral membrane proteins

DeepTMHMM2 was trained to predict RE and IF in TM-α exclusively. However, as the state space of TM-α and soluble proteins is shared (Figure S1), the CRF does not preclude the prediction of such elements in soluble proteins. When analyzing human proteome model predictions, we find that 48 soluble proteins receive an IF or RE. We manually reviewed the plausibility of these predictions (Table S19). Surprisingly, for 24/48 cases, the model’s predictions were supported by either results in literature, or evidence from related proteins that confirmed the predicted element. For another 11 cases, we found evidence that established the protein to be peripheral without elucidating the structural mechanism. For instance, the model predicts an IF in Serine palmitoyltransferase 2 (SPTLC2) that was resolved experimentally by cryo-EM^49^, or in the mitochondrial protein MAIP1, which was experimentally confirmed to be peripherally attached to the membrane^50^ and has its IF confidently predicted by AF2 (Figure 4e).

Among the peripheral cases without direct evidence, we find two IFs in subunit 1 of Dolichol-phosphate mannosyltransferase (DPM1), which is known to be membrane-associated by an undetermined mechanism as part of its enzymatic complex^51^. We also encounter cases where predictions are at odds with annotations in UniProt, such as for the Fatty acid CoA ligase ACSL3, in which the IF is annotated as a TM helix based on sequence analysis, even though experimental results have demonstrated otherwise^52^.

This result allows us to conclude that training on RE and IF regions in TM-α proteins alone can be sufficient to enable the prediction of some modes of peripheral association. However, the number of positive predictions in the human proteome remains low, and we also encounter 6 false positives. In some of these cases, a TM helix of a bitopic protein was erroneously predicted as IF, demonstrating how including RE and IF can render topology prediction more challenging. We therefore see the result as a proof-of-principle that transfer learning from TM proteins is possible, and expect that future general topology methods may achieve more comprehensive coverage by incorporating targeted peripheral training data, overcoming the need for separate peripheral association predictors^24,53^

## Discussion

We introduce DeepTMHMM2, a joint transmembrane topology and membrane type predictor. In addition to handling TM-α, TM-β and SPs like previous methods, DeepTMHMM2 further generalizes the topology prediction objective to also cover TPs, RE and IF.

To overcome the underannotation of RE and IF in available resources, we establish labels by manual review of literature and structural evidence. While absolute performance still lags behind the well-annotated α and β, our curated dataset effectively renders these elements learnable, with the model predicting them correctly in 58.5% and 43.2% of all cases. Presumably, additional data collection efforts may lead to further performance gains.

In rare cases, the model learned to predict RE and IF elements in soluble, non-TM-α proteins. While the prediction of only 48 cases across the human proteome suggests that the recall remains too low to be practically useful, evidence shows that at minimum 50% of the predictions are correct, indicating good precision. While these elements are only sporadically annotated in UniProt as *intramembrane*, we were surprised by the high availability of evidence in primary literature, suggesting an opportunity for targeted data curation for future TM topology models that may include monotopic proteins.

Membrane type prediction further improves the practical utility of topology prediction, ensuring that predictions are directly interpretable for organelles where the generic inside-outside bilayer terminology is ambiguous. However, while prediction works well for cell membranes of different domains and some organelles, other locations such as e.g. the Golgi, still need improvement.

In summary, DeepTHMMM2 offers more comprehensive and interpretable membrane topology predictions. As a sequence-only method, it provides the high throughput expected of a topology predictor, e.g. processing the human proteome in 1 hour and 5 minutes (Methods), making it useful both as a fast first-pass predictor before performing structure predictions, and as a complementary source of information for applications that require both topology and structure information.

## Data and code availability

DeepTMHMM2 is available at https://dtu.biolib.com/DeepTMHMM2.

A Python prediction package is available for download at https://github.com/fteufel/DeepTMHMM2. The training codebase, including all datasets, is available at https://github.com/fteufel/DeepTMHMM2-training.

## Acknowledgements

We thank the teams maintaining PDBTM and OPM for their work providing curated PDB-based topology data. We also thank the authors of TMbed for making cross-validated predictions available for benchmarking.

KDT was supported by NNF grant NNF24SA0100980. OW and AK were in part funded by the Novo Nordisk Foundation through the Center for Basic Machine Learning Research in Life Science (NNF20OC0062606). OW was in part funded by CAZAI (NNF22OC0077058). OW acknowledges support from the Pioneer Center for AI, DNRF grant number P1.

## Online Methods

### Sequence data

We established a dataset for membrane protein topology and membrane type prediction from a collection of sources discussed in detail below. Each sequence *x* of length *L* is annotated with a topology label list *y*^Topology^ and a membrane type label vector *y*^M^. The topology label is of length *L*, with individual position-wise labels *y*^T^ ∈ {*a*, *b*, *α*, *β*, *RE*, *IF*, *SP*, *TP*, *U*} . Soluble regions on the two sides of the membrane are labeled side *a* and side *b*, with side *b* being defined as the side where the N-terminus is exposed after cleavage of a signal peptide, if present, labeled *SP*. Likewise, *TP* denotes N-terminal transit peptide residues. We avoid the more traditional approach of naming the two sides as *inside* and *outside*, as the concept can be ambiguous when also considering organelle membranes. Instead, we assign biological meaning to the two sides via the membrane type label, which defines what *a* and *b* indicate in the specific membrane context (see mapping in Table S4). Transmembrane alpha helices are labeled as *α*, and beta strands as *β*. *RE* and *IF* denote membrane re-entrant and interfacial regions that do not fully traverse the lipid bilayer. *U* is a special label for undetermined residues. We use this at places where the topology is not known due to a lack of experimental evidence or ambiguity. Because the topology label sequence is highly structured (implemented in DeepTMHMM2 by a grammar-based conditional random field, CRF), the possible topology labels for a *U* label at a given position can be quite restrictive. By leveraging the CRF’s capability to *marginalize* over unknown labels, *U* is only used for training and computing evaluation metrics. In inference, DeepTMHMM2 makes biologically valid predictions at all positions, and by design cannot leave any residues undetermined. As each protein can localize to more than one membrane, membrane type labels are multi-hot vectors *y*^M^ ∈ {0,1}^17^ for the 17 membranes (Table S3) modeled by DeepTMHMM2.

Below we describe the collection of the datasets used for training. We have two sources of data: (1) experimental and (2) bootstrapping the experimental with AlphaFold DB and ESM Atlas structures. The experimental set was split at maximum 20% identity by assigning CD-HIT^54^ cluster representatives into five folds, balancing for protein types. Statistics of the partitioned set are given in Table S1.

### SP proteins

Soluble proteins with SPs were sourced from the SignalP 6.0 training dataset by removing any sequence from it that are annotated with a TM region in UniProt, yielding 2,604 sequences. We do not discriminate between the subtypes of SPs. Residues following the SP are labeled as side *b*. For seven proteins without UniProt TM annotations, but ambiguous potential membrane association evidence (such as e.g. pore-forming toxins), *U* was used on mature (non-SP) residues.

### TP proteins

Soluble proteins with TPs were queried from UniProt release 2024_04, requiring experimental evidence for the TP annotation and no additional TM annotations. We include mitochondrial, chloroplast, thylakoid, chromoplast and amyloplast TPs, without discriminating between the subtypes. Fragment proteins are excluded. Any residues following the TP cleavage site (CS) annotation were left undetermined, as we cannot directly discern from a TP annotation whether side *a* or *b* would be applicable.

### Soluble proteins

For soluble proteins, we queried UniProt release 2024_04 for proteins that localize to the cytoplasm (SL-0086), nucleus (SL-0191) and peroxisome (SL-0204) with experimental evidence (ECO:0000269) and a maximum length of 4000. We removed any proteins that

A. are fragments,
B. have an additional transmembrane or intramembrane subcellular location (SL-9904, SL-9905, SL-9906, SL-9907, SL-9908, SL-9909),
C. are annotated with a TM segment, a transit peptide or a signal peptide at any evidence level,
D. Have a secondary location in Mitochondria (SL-0173, SL-0170, SL-0168), plastids (SL-0209), secreted (SL-0243), membrane (SL-0039, SL-0037, SL-0162), and
E. have annotations from high-throughput studies^55,56^ that may reduce label quality, yielding a total of 11,464 sequences.

### Alpha and beta transmembrane proteins

Transmembrane (TM) datasets were compiled using the annotations from the OPM^28^ (version 09 October 2023) and PDBTM^57^ (version 29 November 2024) databases. TM proteins were identified by intersecting entries from both databases, followed by mapping of the shared entries to UniProt using SIFTS^58,59^. Only high-confidence TM segments supported by both resources were retained, requiring an intersection-over-union (IoU) score of at least 0.5 and start/end boundary deviations not exceeding five residues. Since both databases annotate the PDB sequences, we retained UniProt’s annotation regarding signal peptides in the full sequences wherever there was strong evidence for it (ECO:0000269, ECO:0000255, ECO:0000305, ECO:0000250).

We annotated TPs in TM proteins based on UniProt annotations, using the same TP type selection as in the soluble TP protein dataset. In addition to annotating TPs with experimental evidence (ECO:0000269), we also include TPs with weaker evidence levels (ECO:0000303, ECO:0000305, ECO:0000250, ECO:0000255, ECO:0007744) in order to avoid mistakenly labeling the N terminus of a TM protein as *a* or *b* merely because the annotation evidence level is too low (excluding such proteins, as done for the soluble TP set, would be no option, as we wish to preserve these samples because of their experimental TM annotations). However, as we assume that cleavage site annotations may be less accurate without experimental evidence, we do not annotate the full TP for training. Rather, we only label position 1 as TP, followed by undetermined labels *U*, thereby leaving it to the model to learn to predict TPs of suitable length at training time.

RE and IF annotations are available only in PDBTM. However, following discussions with the PDBTM developers (Gábor Tusnády, personal communication), we were informed that these annotations should not be used as training labels in predictive modeling, particularly because of the tendency of the TMDET 3.0 algorithm to predict false positives. Therefore, and to improve data quality, we first performed an initial manual quality control of the regions extracted from PDBTM, followed by extensive manual reannotation of segments in TM-α proteins based on structural and literature evidence. The detailed process is laid out in Supplementary Note 1, and per-protein curation comments are provided in Tables S17-S19. In total, 558 proteins were reannotated with elements, resulting in a total of 540 and 98 proteins being labeled with IF and RE respectively in the final dataset before partitioning.

During manual reannotation, we also curated additional evidence for undetermined regions in alpha and beta TM proteins. Known soluble domains that are not structurally resolved (and therefore labeled as *U*) are labeled as *a* or *b*. Likewise, we annotated SPs and TPs that are mentioned in primary literature, but not recorded in UniProt. For SPs with unknown cleavage site, we use the label in position 1 followed by *U*, as discussed for TPs. All reannotations are documented in Supplementary File 1.

Membrane type annotations were queried from OPM and UniProt release 2024_04, using the mapping defined in Table S5. Conflicts between the two sources were corrected by manual inspection of homologs and original literature. We exclude the OPM membrane types *Peroxisome membrane, Nucleus outer membrane, Cytoplasmic granule membrane* due to a lack of data in the cross-validation set, and *Chloroplast inner membrane* due to random performance (MCC=0.00). *Secreted* and *Gram-positive outer membranes* are excluded due to biological ambiguity. This leaves a total of 17 membrane types for multilabel prediction.

### AlphaFold DB and ESM Metagenomic Atlas transmembrane proteins

We apply a Foldseek-based^31^ pipeline to identify transmembrane proteins with diverse sequences and automatically annotate their topology. The approach is described in detail in Supplementary Note 2, and follows a conservative strategy overall, favoring high label transfer quality over retrieving the maximum number of proteins.

We use our experimental TM-β and TM-α proteins as query sequences to search redundancy-reduced (FoldSeek clustered at a 20% cutoff) AFDB and ESM Atlas databases for proteins with similar predicted structure. After the initial search, we perform an all-vs-all alignment between the found hits and all query proteins and discard hits that align ambiguously with a Template Modeling score >0.5 to seeds with different topologies (e.g. to both a 12TM and 14TM query protein). The remaining pairwise alignments with a Template Modeling score >0.5 and a Foldseek probability >0.99 are used for automated topology label transfer, where we use the Foldseek alignment coordinates to map TM segments from the query protein to the hit protein. As a further alignment quality control, we require that the aligned query TM segment residues have a pLDDT >40 in the hit structure. After transfer, we discard labels/alignments that have incomplete coverage of the TM segments of the query, yield TM segments that are too long or too short given the CRF constraints, or have an average pLDDT <40 over their TM segments. Note that a hit protein can have received multiple labels if it was aligned to more than one query. We aggregate multiple labels per protein into a single consensus label by taking the mean start and end positions of labeled TM segments, discarding hits where TM segments in different labels do not overlap. We homology-reduce the found proteins with respect to the experimental set using MMseqs2 at 20% identity. In total, we obtain 9,660 labeled TM-α and 5,316 labeled TM-β proteins. No membrane type labels are established for these proteins. As we do not have information about orientation, positions in-between the labeled segments are set to undetermined.

### UniProt transmembrane proteins

To increase the available data for learning to predict membrane types, we curated transmembrane proteins with known membrane types but unknown experimental topology from UniProt release 2024_04. We query proteins annotated as transmembrane (either directly as a transmembrane annotation, or locations SL-9905, SL-9906, SL-9907, SL-9908, SL-9909) with any evidence level. We combine the transmembrane status with subcellular location codes and taxonomy filters for the specific membrane type, using manual or experimental evidence codes ECO:0000269, ECO:0000303, ECO:0000305, ECO:0000250. SPARQL queries for each membrane type are provided in the code release. We homology-reduce the found proteins with respect to the experimental set using MMseqs2 at 20% identity, yielding 13,277 proteins. No topology labels are established for these proteins, as the dataset is only used for membrane type prediction.

### The DeepTMHMM2 model

DeepTMHMM2 treats topology prediction as a sequence-to-sequence task, with the input being the protein amino acid sequence, and the output being a residue-wise prediction of segment labels. Multilabel membrane type prediction is performed using an additional classification model. The DeepTMHMM2 model consists of four elements:

A. A protein language model (pLM) embedding of the sequence using ESM-2 (650M),
B. A CNN-LSTM neural network for processing the pLM embeddings,
C. A conditional random field (CRF) decoder for topology prediction, and
D. An attention pooling classification neural network for membrane type prediction given the pLM embeddings.

For each sequence, ESM-2 generates an embedding of size *L* × 1280. We use the embeddings as frozen input features and do not fine-tune ESM-2 during training. Like in SignalP 5.0, we first apply an initial convolution layer with kernel size 3 before processing the sequence with the LSTM. The LSTM outputs hidden states of size *L* × ℎ, where ℎ = 512 is the output size of both the CNN and the LSTM. We perform a linear projection of ℎ to the label dimension, obtaining a matrix of unnormalized scores (CRF emissions) *L* × *C*, where *C* is the number of states.

The (linear-chain) CRF^26^ is an undirected graphical probabilistic model that models sequences as paths through a predefined state space. Rather than treating the sequence-to-sequence problem as independent predictions at each position, the CRF computes a joint distribution over the full sequence using emissions *ψ* and transitions *φ* for the modeled states. The emissions are unnormalized position-wise state probabilities of size *L* × *C*. The transitions are a matrix of size *C* × *C* that encodes the unnormalized probabilities of transitioning between states on adjacent sequence positions.

We design the state space to explicitly enforce the known biophysical structure of the prediction problem. We refer to this known structure (a set of rules for forming syntactically valid topology strings) as grammar. For instance, it is impossible for a valid topology to jump from side *a* of the membrane to side *b*, without there being a transmembrane segment of sufficient length in-between. Likewise, a re-entrant region or interfacial helix starting on side *a* cannot reemerge on side *b*. In summary, we enforce the following biophysical constraints in the model:

● The membrane can only be traversed by having a complete transmembrane segment.
● Transmembrane segments need to end on the opposite membrane side from where they started and have a minimum and maximum length.
● Interfacial and reentrant segments need to start and end on the same side of the membrane and have a minimum length.
● SPs and TPs, also have a minimum length, are always at the N terminus, and SPs end in membrane side *b*.
● A protein cannot contain TM helices and beta sheets simultaneously.

To encode this grammar in the state space, we make use of an expanded number of states that allows the CRF to track the length and orientation of segments. For the orientation, we “split” the states for the membrane segments {*α*, *β*, *RE*, *IF*} into two, so that we obtain the orientated segments {*α^ab^*, *α^ba^*, *β^ab^*,*β^ba^*, *RE^aa^*,*RE^bb^*, *IF^aa^*, *IF^bb^*}. To model the length constraints, like in the original TMHMM model^60^, we represent each segment as multiple states up to the maximum length (*α*: 30, *β*: 15), so that we obtain e.g. the state sequence [*α*_1_*^ab^*, *α*_2_*^ab^*, *α*_3_*^ab^*, *α*_4_*^ab^*. . . *α*_29_*^ab^*, *α*_30_*^ab^*] for *α^ab^* segments.

Segments without a defined maximum length follow the same approach for the first states until reaching a minimum length, followed by a state with a self-loop. Moreover, we split the two membrane sides into *a_α_*, *b_α_*, *a_β_*, *b_β_* to prevent spurious mixing of segments. This yields a space with 135 (60 *α* + 30 *β*+ 10 *SP* + 5 *TP* + 16 *RE* + 10 *IF* + 2 *a* + 2 *b*) states in total.

In this expanded state space, we enforce the grammar by constraining the CRF’s transitions. For instance, in the *α^ab^* segment, only membrane side state *a_α_* can transition to *α^ab^*. The initial states up to half the minimum length (*α^ab^*) transition deterministically to the following state of the same segment only. To enable variable length, state *α*_5_*^ab^* is allowed to “skip ahead” and directly transition into any of [*α*_6_*^ab^*, . . ., *α*_26_*^ab^*]. The final states up to *α^ab^* again are only transitioning to their following state, thereby (together with the initial states) enforcing the minimum length constraint. *α*_30_*^ab^* can only transition to membrane side *b_α_*. The same state construction scheme is applied to all segments. States *a_α_*, *b_α_*, *a_β_*, *b_β_* are allowed to self-transition indefinitely to model soluble domains of any length. The complete state space with all transitions is given in Supplementary Figure 1. Constraining is performed by clamping all disallowed transitions to zero. The values of the valid transitions are learned end-to-end during training.

By default, this state space would imply that our neural network needs to predict emissions for 135 distinct states. However, as our state space is highly redundant from a biological feature perspective and only serves the purpose of enforcing the grammar, we only predict 10 emissions for {*α*, *β*, *RE*, *IF*, *SP*, *TP*, *a_α_*, *b_α_*, *a_β_*, *b_β_*}, and share these between all states of a given segment by repeating the *L* × 10 neural network output to size *L* × 135. CRF marginal likelihoods are computed using the forward-backward algorithm, and the most likely topology sequence *y*D*^T^* is decoded using the Viterbi algorithm. After prediction, we map the states back to the original label alphabet {*a*, *b*, *α*, *β*, *RE*, *IF*, *SP*, *TP*}. The predicted type of the protein is inferred by simply evaluating the presence of segment labels in the predicted topology sequence (e.g. if a predicted topology sequence contains at least one *α* label, the predicted type will be *α*).

The membrane type is predicted by a lightweight classifier based on the DeepLoc 2.0^27^ architecture for multilabel prediction. It takes the ESM embeddings as input and performs attention pooling before feeding the representation to an MLP for classification.

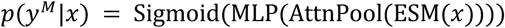

To convert predicted probabilities into multilabel predictions, we apply class-specific decision thresholds that are calibrated on the validation set. When returning a topology prediction, the membrane type with the highest predicted probability is used to map the labels *a*, *b* to biological terms, given the mapping in Table S4.

### Training

We make use of the multi-tag CRF framework to train on partially labeled topologies containing *U*. This allows us to sum the likelihood over all CRF states that a *U* could represent at a given position. To maximize the information content provided by undetermined positions, we decide which states to set to true based on the protein’s type and determined positions using a collection of general rules. For any contiguous segment of *U* labels, we set the following states to true in the multi-tag label matrix of size *L* × *C*:

1. Non-membrane states: *a* and *b* are always true at all positions
2. If the unknown segment is at the N-terminus of the sequence, all possible *SP* and *TP* configurations that fit the segment’s length are set to true
3. Alpha-helical proteins: All possible *α*, *RE*, *IF* configurations that fit the segment’s length are set to true
4. Beta-barrel proteins: All possible *β* configurations are set to true, unless the protein already contains at least 8 strands, in which case it is assumed to be a full barrel that cannot permit any additional TM strand.

Additionally, depending on the available evidence for an unknown segment, the rule set can be further restricted to only allow subsets of labels. In the dataset files, we use the letter ”u” to denote regions determined to be restricted to non-membrane spanning and non-signal segments (permitting only *a*, *b*, *RE*, *IF* predictions), and ”.” for N-terminal undetermined regions that additionally permit an unknown sorting signal (*a*, *b*, *RE*, *IF*, *TP*, *SP*). The generic ”U” remains unrestricted, allowing all prediction types.

Given the collected true states at each position *l* < *L*, let *Y* be the set of all true paths through the state space. The log likelihood is then computed as

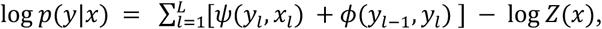

with *Z*(*x*) being the CRF’s partition function:

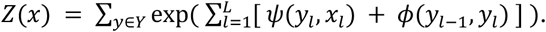

Due to the nearest neighbour coupling, the summation over *Y* in the partition function can be evaluated linear in *L* using the forward algorithm.

The topology prediction model is trained for two epochs, training five models in cross-validation on the experimental set split. The AFDB/ESMAtlas derived dataset can be mixed in for all folds, as sequences are already always reduced to 20% identity to the experimental data and therefore do not induce leakage at this threshold. We explore the hyperparameter space of the model in cross-validation using Optuna^61^ by varying the data composition of training batches as well as the learning and dropout rate of the model. As the criterion, validation metrics are summarized as the mean of 0.6 * topology prediction performance (sum of topology % correct) and 0.4 * type prediction performance (sum of F1 scores).

The membrane type classifier is trained separately from the topology prediction network for five epochs. For multilabel membrane type prediction, we optimize the binary cross-entropy given the label vector *y^M^*. We train with five-fold cross-validation on the experimental data set, using the same split assignments as for topology prediction. Like the AFDB/ESMAtlas dataset, the UniProt dataset can be mixed in for all folds as it was reduced at 20%. We train with batches of 24 experimental + 24 UniProt samples. The model uses a hidden dimension of 512 and is trained for five epochs at a learning rate of 1e-4. After training, we determine an optimal decision threshold for each class by maximizing the Matthews Correlation Coefficient (MCC) on the validation set.

### Metrics

Given a true and predicted topology of a protein, we compute multiple binary correctness criteria that together assess the prediction of all TM segments, the protein’s membrane orientation, any signal or transit peptide, as well as re-entrant and interfacial segments.

- Transmembrane segments count as predicted correctly if all TM segments are predicted with a minimal overlap of 5 (alpha) or 3 (beta) residues.
- For the protein’s orientation, we check that the predicted side of the residue preceding each TM segment matches the truth (this sufficiently implies that the TM segment is correctly orientated).
- Re-entrant and interfacial segments are evaluated using a 5-residue minimal overlap.
- Signal and target peptides are evaluated at tolerance windows (0-3 residues) around their true cleavage site.
- Sections of the protein that are labeled as undetermined are ignored when computing the criteria, thereby allowing for the prediction of unlabeled features without affecting any metrics.

Matching previous work, we combine these criteria into a single topology correctness measure for alpha and beta TM proteins. A TM protein topology is considered correct if the TM segments are correct, the orientation is correct, and, when applicable, the signal peptide is correct (at tolerance window 3). Transit peptides, re-entrant and interfacial elements are omitted from the combined criterion to enable benchmarking against other methods that do not predict these elements.

As the metric for topology prediction, we compute the % correct. For protein type prediction, we compute the F1 score, using all other types as the negative class. When determining the predicted type, additional predicted segments that start or end in undetermined sections are ignored for computing false positives (E.g., a protein that does not have any *RE* in its true label but has an RE feature predicted in an undetermined section does not count as false positive for type RE).

For membrane type prediction performance, we report the MCC, F1, AUC and AUPRC of each class. Standard errors for accuracy and percentage correctness were computed using the binomial standard error. Standard errors for F1 scores and MCCs were estimated using the delta method based upon the multinomial distribution over confusion matrix entries^62^.

### Benchmarking

We perform three benchmark experiments, comparing performance to TMbed, DeepTMHMM1 and SignalP 6.0. For these methods, cross-validated predictions are available, enabling direct comparison without the risk of data leakage. For each method, we subset the available cross-validated predictions to the proteins that are common with the DeepTMHMM2 cross-validation set, yielding three benchmark datasets of 3287, 1380, and 4902 proteins. For TMbed, we additionally filter proteins that were blacklisted in the available material due to potential leakage.

Additionally, we benchmark a range of popular topology prediction methods in their publicly available implementations, using a recent review as reference^63^. We focus on methods that perform complete topology prediction (with or without SPs), excluding methods that only annotate TM segment positions without resolving orientation. Likewise, methods whose official web server is offline were excluded. A full list of methods is provided in Table S10. We note that in this regime, train set data leakage cannot be avoided, leading to potential overestimation of real-world performance against cross-validated DeepTMHMM2 predictions. Full benchmarking experiments are in Tables S11-S15. Beyond sequence-based topology prediction methods, we also investigate the potential performance of structure-based positioning and segment detection algorithms when applied to AlphaFold 2 predictions in Supplementary Note 3.

### Ensemble prediction

The final DeepTMHMM2 predictor is an ensemble of the cross-validated models for topology prediction and membrane type prediction. Membrane type predictions are ensembled by averaging the predicted probabilities of the five models.

As topologies are discrete and cannot be averaged directly, we perform aggregation by choosing the prediction with the highest agreement between the five models. For each of the five predicted topologies, we count how many other predicted topologies are equivalent by evaluating matching sorting signal presence and membrane feature overlap. We return the topology with the highest agreement count and its corresponding marginal probabilities. Running on an RTX 8000 GPU, the predictor takes approximately 1 hour and 5 minutes to process the UniProt human proteome (excluding Titin, which at a length of more than 34,000 AAs cannot be processed on an RTX 8000).

### Membrane biology analysis of predictions

We group the cross-validated TM-α predictions by their true membrane type. Proteins labeled with multiple membrane types contribute to the summary statistics of each labeled membrane type for all analyses. For the aromatic belt analysis, we align all TM-α segments at their N-terminal and C-terminal boundaries and compute the position-wise frequency of Tryptophan (W) and Tyrosine (Y) within -10 to +10 residues of the boundary. Positions on the soluble side are only included if all intervening residues between that position and the TM boundary are themselves soluble, ensuring adjacent TM regions are excluded when the soluble loop is shorter than 10.

For the positive-inside rule analysis, for each protein, we collect all predicted soluble loop regions that fall within 10 residues of any TM segment boundary. This is done so that the analysis stays focused on flanking soluble loops, omitting larger soluble domains. We count all occurrences of Lysine (K) and Arginine (R) residues in loops on side a and loops on side b. The K+R fraction is computed as the K+R count of side a, divided by the total K+R count. Proteins for which neither side carries any K or R near a membrane boundary are excluded. A value above 0.5 therefore indicates enrichment of positively charged residues on the side that is topologically understood as cytoplasmic/inside.

### Gene Ontology enrichment analysis of RE and IF proteins

We obtained DeepTMHMM2 ensemble predictions for all proteins in Swiss-Prot and filtered for those predicted to have at least one TM-α segment. Using Goatools^64^, we conducted GO enrichment analysis for Molecular Function terms using Fisher’s exact test. The background population was defined as all predicted TM-α proteins from Swiss-Prot possessing at least one Molecular Function annotation (n=61,933). We compared proteins containing RE or IF features against this background, correcting p-values for multiple testing via the Benjamini-Hochberg method (adjusted $p < 0.05$). The analysis restricted term selection to a fixed DAG depth of five. To further reduce redundancy on this depth level, descendants of the terms ’G protein-coupled receptor activity’ (GO:0004930), ’monoatomic ion transmembrane transporter activity’ (GO:0015075), ’transferase activity’ (GO:0016740), ’chlorophyll binding’ (GO:0016168), and ’oxidoreductase activity’ (GO:0016491) were manually pruned from the graph. Prevalence of GO terms within the feature-positive groups was quantified by the gene ratio, defined as the fraction of proteins possessing the specific feature (RE/IF) that are annotated with a given GO term. Enrichment of GO terms against the background distribution was calculated as the fold change of the gene ratio over the background rate.

### Topology-specific enrichment of ClinVar pathogenic variants

We obtained DeepTMHMM2 ensemble predictions for the UniProt reference human proteome and filtered for proteins predicted to have at least one TM-α segment with at least one ClinVar variant. Each ClinVar variant (GRCh38) classified as Pathogenic or Likely Pathogenic (collectively Pathogenic) or Benign or Likely Benign (collectively Benign) was mapped to its corresponding protein via the UniProt gene symbol and assigned to one of the four topological categories TM-α, RE, IF, or soluble regions. For variants spanning multiple residues, assignment followed the highest-priority overlapping feature (RE > IF > TM-α > soluble).

To account for differences in the total number of residues assigned to each topological category, we modeled variant counts using a Poisson generalized linear model with a log link and the logarithm of total residues per category as an offset term. The model included topological category, variant class (pathogenic versus benign), and their interaction as explanatory variables.

The resulting interaction term estimates the length-normalized relative rate ratio (RR) of pathogenic to benign variants in each feature relative to soluble regions as the baseline, with 95% confidence intervals and p-values derived directly from the GLM.

## Supplementary Material

- **Supplementary File 1**

- Supplementary Note 1: Refinement of transmembrane protein annotations
- Supplementary Note 2: Mining structure databases for transmembrane proteins
- Supplementary Note 3: Evaluating TMDET and PPM
- Figure S1. CRF state space.
- Table S1. Summary statistics per cross-validation fold of the DeepTMHMM2 experimental transmembrane topology dataset.
- Table S2. Membrane type label counts of the experimental transmembrane topology dataset
- Table S3. Label counts of the UniProt membrane type multilabel dataset.
- Table S4.
- Mapping of the two modeled topological sides A and B to specific locations based on membrane type.
- Table S5: Mapping of UniProt locations to OPM membrane types.
- Table S6. Metrics on the full cross-validated dataset.
- Table S7. Classification metrics on the TMbed cross-validated benchmark dataset.
- Table S8. Classification metrics on the DeepTMHMM 1 cross-validated benchmark dataset.
- Table S9. Classification metrics on the SignalP cross-validated benchmark dataset.
- Table S10. The full list of methods hat were used for the benchmark.
- Table S11: Extended benchmark for TM-ɑ prediction performance.
- Table S12: Extended benchmark for TM-β prediction performance.
- Table S13: Extended benchmark for RE prediction performance.
- Table S14: Extended benchmark for IF prediction performance.
- Table S15: Extended benchmark for PRED-TMSdeep.
- Table S16: Membrane type prediction performance.
- Figure S2. % correct for SP prediction at varying tolerance windows around the true cleavage site
- Figure S3. % correct for TP prediction at varying tolerance windows around the true cleavage site
- Supplementary File 2

- Table S17: Experimental set manual topology curation
- Table S18: Unresolved region manual curation in TM-ɑ proteins
- Table S19: Unresolved region manual curation in TM-β proteins
- Table S20: Manual review of predicted RE/IF segments in soluble proteins

## Bibliography

1. Tsirigos, K. D. et al. Topology of membrane proteins-predictions, limitations and variations. Curr. Opin. Struct. Biol. 50, 9–17 (2018).

2. Gong, J. et al. Understanding Membrane Protein Drug Targets in Computational Perspective. Curr. Drug Targets 20, 551–564 (2019).

3. Hendrickson, W. A. Atomic-level analysis of membrane-protein structure. Nat. Struct. Mol. Biol. 23, 464–467 (2016).

4. Cheng, Y. Single-Particle Cryo-EM at Crystallographic Resolution. Cell 161, 450–457 (2015).

5. Jumper, J. et al. Highly accurate protein structure prediction with AlphaFold. Nature 596, 583– 589 (2021).

6. Abramson, J. et al. Accurate structure prediction of biomolecular interactions with AlphaFold 3. Nature 630, 493–500 (2024).

7. Hegedűs, T., Geisler, M., Lukács, G. L. & Farkas, B. Ins and outs of AlphaFold2 transmembrane protein structure predictions. Cell. Mol. Life Sci. CMLS 79, 73 (2022).

8. Swanson, E., Nichols, M., Ravichandran, S. & Ogden, P. mBER: Controllable de novo antibody design with million-scale experimental screening. 2025.09.26.678877 Preprint at 10.1101/2025.09.26.678877 (2025).

9. Soh, T. K. et al. A proteome-wide structural systems approach reveals insights into protein families of all human herpesviruses. Nat. Commun. 15, 10230 (2024).

10. Ferguson, L. et al. DeorphaNN: Virtual screening of GPCR peptide agonists using AlphaFold-predicted active-state complexes and deep learning embeddings. 2025.03.19.644234 Preprint at 10.1101/2025.03.19.644234 (2025).

11. Teufel, F. et al. Deorphanizing Peptides Using Structure Prediction. J. Chem. Inf. Model. 63, 2651–2655 (2023).

12. von Heijne, G. Membrane protein structure prediction. Hydrophobicity analysis and the positive-inside rule. J. Mol. Biol. 225, 487–494 (1992).

13. Claros, M. G. & von Heijne, G. TopPred II: an improved software for membrane protein structure predictions. Comput. Appl. Biosci. CABIOS 10, 685–686 (1994).

14. Krogh, A., Larsson, B., von Heijne, G. & Sonnhammer, E. L. Predicting transmembrane protein topology with a hidden Markov model: application to complete genomes. J. Mol. Biol. 305, 567–580 (2001).

15. Sonnhammer, E. L., von Heijne, G. & Krogh, A. A hidden Markov model for predicting transmembrane helices in protein sequences. Proc. Int. Conf. Intell. Syst. Mol. Biol. 6, 175–182 (1998).

16. Bathla, D., Mishra, R. & Ahmad, S. A Survey of Current Status in AI-Based Topology Prediction of Transmembrane Proteins. Methods Mol. Biol. 2947, 109–135 (2025).

17. Tsirigos, K. D., Peters, C., Shu, N., Käll, L. & Elofsson, A. The TOPCONS web server for consensus prediction of membrane protein topology and signal peptides. Nucleic Acids Res. 43, W401–407 (2015).

18. Bagos, P. G., Liakopoulos, T. D. & Hamodrakas, S. J. Evaluation of methods for predicting the topology of beta-barrel outer membrane proteins and a consensus prediction method. BMC Bioinformatics 6, 7 (2005).

19. Dobson, L., Reményi, I. & Tusnády, G. E. CCTOP: a Consensus Constrained TOPology prediction web server. Nucleic Acids Res. 43, W408–412 (2015).

20. Hallgren, J. et al. DeepTMHMM predicts alpha and beta transmembrane proteins using deep neural networks. 2022.04.08.487609 Preprint at 10.1101/2022.04.08.487609 (2022).

21. Bernhofer, M. & Rost, B. TMbed: transmembrane proteins predicted through language model embeddings. BMC Bioinformatics 23, 326 (2022).

22. Viklund, H. & Elofsson, A. OCTOPUS: improving topology prediction by two-track ANN-based preference scores and an extended topological grammar. Bioinformatics 24, 1662–1668 (2008).

23. Nugent, T. & Jones, D. T. Transmembrane protein topology prediction using support vector machines. BMC Bioinformatics 10, 159 (2009).

24. Feng, S.-H., Xia, C.-Q., Zhang, P.-D. & Shen, H.-B. Ab-Initio Membrane Protein Amphipathic Helix Structure Prediction Using Deep Neural Networks. IEEE/ACM Trans. Comput. Biol. Bioinform. 19, 795– 805 (2022).

25. Lin, Z. et al. Evolutionary-scale prediction of atomic-level protein structure with a language model. Science 379, 1123–1130 (2023).

26. Lafferty, J. D., McCallum, A. & Pereira, F. C. N. Conditional Random Fields: Probabilistic Models for Segmenting and Labeling Sequence Data. in *Proceedings of the Eighteenth International Conference on Machine Learning* 282–289 (Morgan Kaufmann Publishers Inc., San Francisco, CA, USA, 2001).

27. Thumuluri, V., Almagro Armenteros, J. J., Johansen, A. R., Nielsen, H. & Winther, O. DeepLoc 2.0: multi-label subcellular localization prediction using protein language models. Nucleic Acids Res. 50, W228–W234 (2022).

28. Lomize, M. A., Pogozheva, I. D., Joo, H., Mosberg, H. I. & Lomize, A. L. OPM database and PPM web server: resources for positioning of proteins in membranes. Nucleic Acids Res. 40, D370–376 (2012).

29. Dobson, L. et al. UniTmp: unified resources for transmembrane proteins. Nucleic Acids Res. 52, D572–D578 (2024).

30. The UniProt Consortium. UniProt: the Universal Protein Knowledgebase in 2025. Nucleic Acids Res. 53, D609–D617 (2025).

31. van Kempen, M. et al. Fast and accurate protein structure search with Foldseek. Nat. Biotechnol. 42, 243–246 (2024).

32. Varadi, M. et al. AlphaFold Protein Structure Database in 2024: providing structure coverage for over 214 million protein sequences. Nucleic Acids Res. 52, D368–D375 (2023).

33. Almagro Armenteros, J. J., et al. Detecting sequence signals in targeting peptides using deep learning. Life Sci. Alliance 2, e201900429 (2019).

34. Sharpe, H. J., Stevens, T. J. & Munro, S. A Comprehensive Comparison of Transmembrane Domains Reveals Organelle-Specific Properties. Cell 142, 158–169 (2010).

35. van Meer, G., Voelker, D. R. & Feigenson, G. W. Membrane lipids: where they are and how they behave. Nat. Rev. Mol. Cell Biol. 9, 112–124 (2008).

36. Bretscher, M. S. & Munro, S. Cholesterol and the Golgi Apparatus. Science 261, 1280–1281 (1993).

37. Glushkova, D., Böhm, S. & Beck, M. Systematic membrane thickness variation across cellular organelles revealed by cryo-ET. J. Cell Biol. 225, e202504053 (2025).

38. Killian, J. A. & von Heijne, G. How proteins adapt to a membrane-water interface. Trends Biochem. Sci. 25, 429–434 (2000).

39. Landolt-Marticorena, C., Williams, K. A., Deber, C. M. & Reithmeier, R. A. Non-random distribution of amino acids in the transmembrane segments of human type I single span membrane proteins. J. Mol. Biol. 229, 602–608 (1993).

40. Yau, W. M., Wimley, W. C., Gawrisch, K. & White, S. H. The preference of tryptophan for membrane interfaces. Biochemistry 37, 14713–14718 (1998).

41. Heijne, G. The distribution of positively charged residues in bacterial inner membrane proteins correlates with the trans-membrane topology. EMBO J. 5, 3021–3027 (1986).

42. von Heijne, G. & Gavel, Y. Topogenic signals in integral membrane proteins. Eur. J. Biochem. 174, 671–678 (1988).

43. Wallin, E. & von Heijne, G. Genome-wide analysis of integral membrane proteins from eubacterial, archaean, and eukaryotic organisms. Protein Sci. Publ. Protein Soc. 7, 1029–1038 (1998).

44. von Heijne, G. Membrane protein structure prediction: Hydrophobicity analysis and the positive-inside rule. J. Mol. Biol. 225, 487–494 (1992).

45. Delaney, E., Khanna, P., Tu, L., Robinson, J. M. & Deutsch, C. Determinants of pore folding in potassium channel biogenesis. Proc. Natl. Acad. Sci. U. S. A. 111, 4620–4625 (2014).

46. Huynh, J., Thomas, W. G., Aguilar, M.-I. & Pattenden, L. K. Role of helix 8 in G protein-coupled receptors based on structure–function studies on the type 1 angiotensin receptor. Mol. Cell. Endocrinol. 302, 118–127 (2009).

47. Landrum, M. J. et al. ClinVar: public archive of relationships among sequence variation and human phenotype. Nucleic Acids Res. 42, D980–D985 (2014).

48. Frazer, J. et al. Disease variant prediction with deep generative models of evolutionary data. Nature 599, 91–95 (2021).

49. Wang, Y. et al. Structural insights into the regulation of human serine palmitoyltransferase complexes. Nat. Struct. Mol. Biol. 28, 240–248 (2021).

50. König, T. et al. The m-AAA Protease Associated with Neurodegeneration Limits MCU Activity in Mitochondria. Mol. Cell 64, 148–162 (2016).

51. Maeda, Y., Tanaka, S., Hino, J., Kangawa, K. & Kinoshita, T. Human dolichol-phosphate-mannose synthase consists of three subunits, DPM1, DPM2 and DPM3. EMBO J. 19, 2475–2482 (2000).

52. Poppelreuther, M. et al. The N-terminal region of acyl-CoA synthetase 3 is essential for both the localization on lipid droplets and the function in fatty acid uptake. J. Lipid Res. 53, 888–900 (2012).

53. Sapay, N., Guermeur, Y. & Deléage, G. Prediction of amphipathic in-plane membrane anchors in monotopic proteins using a SVM classifier. BMC Bioinformatics 7, 255 (2006).

54. Li, W. & Godzik, A. Cd-hit: a fast program for clustering and comparing large sets of protein or nucleotide sequences. Bioinformatics 22, 1658–1659 (2006).

55. Ding, D. Q., et al. Large-scale screening of intracellular protein localization in living fission yeast cells by the use of a GFP-fusion genomic DNA library. Genes Cells Devoted Mol. Cell. Mech. 5, 169–190 (2000).

56. Matsuyama, A., et al. ORFeome cloning and global analysis of protein localization in the fission yeast Schizosaccharomyces pombe. Nat. Biotechnol. 24, 841–847 (2006).

57. Kozma, D., Simon, I. & Tusnády, G. E. PDBTM: Protein Data Bank of transmembrane proteins after 8 years. Nucleic Acids Res. 41, D524–529 (2013).

58. Velankar, S., et al. SIFTS: Structure Integration with Function, Taxonomy and Sequences resource. Nucleic Acids Res. 41, D483–489 (2013).

59. Dana, J. M., et al. SIFTS: updated Structure Integration with Function, Taxonomy and Sequences resource allows 40-fold increase in coverage of structure-based annotations for proteins. Nucleic Acids Res. 47, D482–D489 (2019).

60. Krogh, A., Larsson, B., von Heijne, G. & Sonnhammer, E. L. Predicting transmembrane protein topology with a hidden Markov model: application to complete genomes. J. Mol. Biol. 305, 567–580 (2001).

61. Akiba, T., Sano, S., Yanase, T., Ohta, T. & Koyama, M. Optuna: A Next-generation Hyperparameter Optimization Framework. in Proceedings of the 25th ACM SIGKDD International Conference on Knowledge Discovery & Data Mining 2623–2631 (Association for Computing Machinery, New York, NY, USA, 2019). doi:10.1145/3292500.3330701.

62. Casella, G. & Berger, R. Statistical Inference. (Chapman and Hall/CRC, New York, 2024). doi:10.1201/9781003456285.

63. Bathla, D., Mishra, R. & Ahmad, S. A Survey of Current Status in AI-Based Topology Prediction of Transmembrane Proteins. Methods Mol. Biol. 2947, 109–135 (2025).

64. Klopfenstein, D. V., et al. GOATOOLS: A Python library for Gene Ontology analyses. Sci. Rep. 8, 10872 (2018).

