## Supplementary Information for "DeepTMHMM2 enables accurate prediction of transmembrane protein topology and subcellular location"

### Supplementary Note 1: Refinement of transmembrane protein annotations

In initial training runs on a preliminary dataset with reentrant regions (RE) and interfacial helices (IF) sourced from PDBTM, we observed that while the model was able to learn these patterns from the training data to some extent, performance on held out data was low and overly sensitive to different training data splits. Manual inspection of the PDBTM annotations showed that, in many cases, putative segments of interest in the 3D structure were missed and mislabeled as soluble, even though the membrane insertion by TMDET[1] correctly positioned the segment with respect to the lipid bilayer, the segments were successfully detected by TMDET in very close homologs, and the structure's original publication described the segment in question correctly.

We believe that this is due to the segment detection methods[2] being overly sensitive to slight mispositioning of the bilayer: For TM segments, any shift of the bilayer along the Z axis (the computed membrane normal) would leave the segment annotation mostly correct, merely shifting the inferred start and end of the segment by a few residues. However, the same shift could prevent an interfacial or re-entrant segment from being called, as the method imposes strict requirements on its relative positioning with respect to the predicted plane. This threshold sensitivity is understood to be a critical factor, with the recent version 4.0 of TMDET[3] now exposing its detection parameters to the user to enable adjustment on a case-by-case basis for specific proteins.

Therefore, we opted to manually refine topology annotations based on structural evidence not detected or not considered by PDBTM. We started our curation by first manually inspecting all existing RE and IF annotations in the proteins of the TM-α set, considering supporting evidence in original papers, raw structures (resolved lipids, amphipathicity) and known features of protein families. Dubious annotations were discarded, and additionally identified segments annotated or corrected based on found evidence.

Next, we annotated alpha-TM proteins without labeled RE/IF segments. To make inspection tractable and prioritize candidate proteins for reannotation, we identified putative RE/IF segments using

1. known features of protein families (e.g. missing annotations of helix 8 in GPCRs[4] or D/E helices in LHC proteins[5] could be identified at large scale),
2. disagreements between homologs revealed by MMseqs2[6] sequence clustering, and
3. model predictions using DeepTMHMM2 models trained on preliminary data.

It is important to note that also for approach 3, preliminary predictions are never used for annotation decisions - the predictor only serves as a prioritization helper. Segments were then annotated based on supporting literature statements, comparison with homologs and alternative structures of the same protein, and resolved lipids in the structure that lend further support to inferred bilayer planes. We perform multiple iterations of this procedure, with the dataset used for finding disagreements and training preliminary models gradually improving. For each annotated protein, we provide a curation note in Table S16.

For some proteins that have structurally ambiguous putative RE/IF segments, we were not able to curate sufficient evidence (e.g. PsaA and PsaB of Photosystem I) to support reannotation. To avoid introducing a false negative training signal, soluble regions were set to undetermined in such cases.

### Supplementary Note 2: Mining structure databases for transmembrane proteins

We use our annotated experimental dataset as the starting point for identifying transmembrane proteins with highly diverse amino acid sequences but conserved structure. We apply Foldseek[7] to search the predicted structure databases AlphaFold DB[8] and ESM Atlas[9] for proteins with structural homology. As we are only interested in proteins with less than 20% identity to any other annotated protein due to our homology reduction strategy applied for cross-validation, we first reduce the Foldseek databases AFDB50 and ESM30 to 20% identity using Foldseek's clustering module. On the reduced databases, we perform a Foldseek search with default parameters for each annotated experimental sequence, limiting the number of returned hits to 100,000 each. To avoid alignment to soluble domains, we trim the AlphaFold structures of experimental sequences to their annotated TM regions. Single-pass TM proteins are omitted, as their trimmed structures reduce to a single helix, providing limited structural context for homology searches. We keep the 5 preceding and following amino acids as well as all intermittent loops between TM segments.

To further reduce the number of hits we perform a first homology reduction of the hits using MMseqs2 clustering at 20%. When reducing, AlphaFold DB hits have retention priority over ESM Atlas hits. The following steps serve to ensure that hits are annotated with TM regions at high quality. We perform an all-vs-all Foldseek alignment of the experimental seed structures to the hit structures, discarding alignments below a TM (Template Modeling)-score[10] of 0.5. The remaining alignments are used for label transfer and quality control. If a hit was aligned to multiple experimental sequences that have divergent transmembrane topology (e.g. a hit has alignments to both a 6TM and a 7TM protein with TM-score>0.5), the hit is discarded. We further filter using FoldSeek's hit probability, removing alignments below 99% probability. For the remaining alignments, we create a transmembrane topology annotation by transferring the experimental annotation according to the aligned residues. If the transferred annotation has less TM segments than the experimental annotation, the alignment is discarded. We also remove alignments that contain TM segments that do not follow the respective minimum and maximum length for helix (10-30) or beta (5-15) segments. Hits that have an average pLDDT lower than 70 in their transmembrane residues are discarded.

### Supplementary Note 3: Evaluating TMDET and PPM on AlphaFold predicted structures without curation

Given the increasing feasibility of large-scale protein structure prediction and the availability of predicted structures in databases, structure-based identification of membrane proteins and the delineation of their transmembrane regions are increasingly important. The PDBTM and OPM databases do this on experimental structures by applying the physics-based TMDET and PPM algorithms, followed by additional manual curation and refinement of detected topologies. However, when the aim is to perform large-scale topology inference on predicted structures, such manual refinement would become infeasible. To assess the performance of a potential direct application of such algorithms, we applied TMDET 4.0[3] and PPM 2.0[11] directly to the AlphaFold predicted structures of our experimental dataset (Tables S11-S14). Because these methods do not predict topology orientation, evaluation was restricted to segment overlap for α-helical and β-barrel proteins and, in the case of TMDET, to re-entrant (RE) and interfacial (IF) regions as well (PPM 2.0 does not annotate them).

AF2+PPM achieved segment-level F1 scores of 0.90 for both α-helical and β-barrel proteins, whereas AF2+TMDET reached F1 scores of 0.65 and 0.84, respectively. However, both structure-based approaches showed substantially lower topology accuracy than DeepTMHMM2. The largest differences were observed for membrane-associated structural elements beyond canonical transmembrane segments. For re-entrant regions, DeepTMHMM2 achieved an F1 score of 0.72 and correctly identified 58.5% of proteins, compared with 0.39 and 51.2% for AF2+TMDET. Performance differences were even more pronounced for interfacial helices, where DeepTMHMM2 attained an F1 score of 0.73 and 43.3% correct topologies, whereas AF2+TMDET reached an F1 score of 0.38 and only 15.3% correct predictions. Our results therefore show that direct application of physics-based insertion underperforms sequence-based approaches, motivating the use of predictions before application[12]. Thus, DeepTMHMM2 not only identifies membrane proteins and their transmembrane segments directly from sequence, but also provides substantially improved characterization of more complex membrane-associated topologies without requiring structural models or additional post-processing.

### Supplementary Figures and Tables

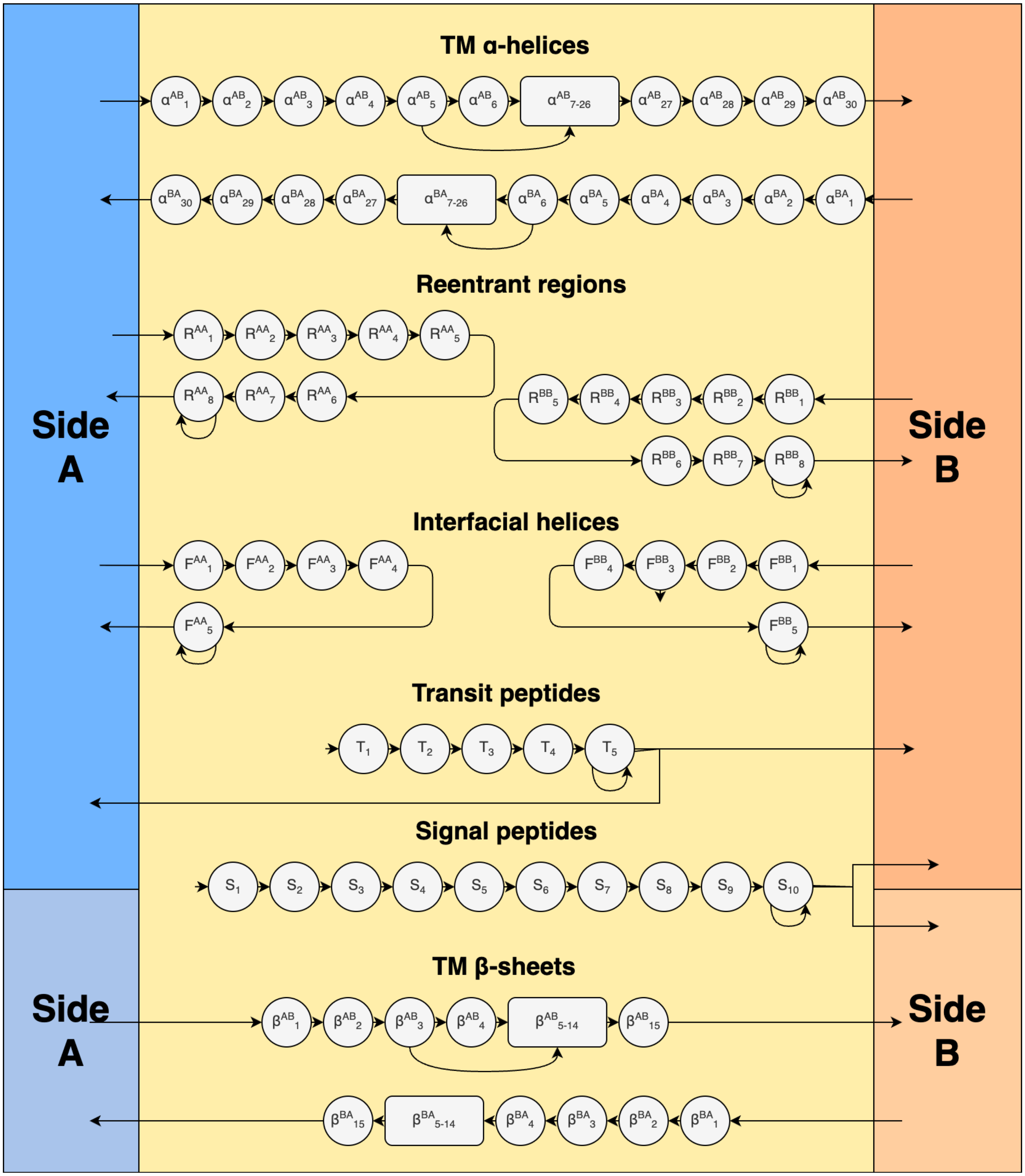


**Figure S1.** The state space of the CRF. Circles represent individual states, while rectangles indicate groups of states. Arrows denote allowed state transitions. Topologies start in $\{a, b, S_1, T_1\}$, and end in $\{a, b\}$. The side states are split for alpha helical and beta barrel proteins, with no transitions allowed.

**Table S1.** Summary statistics per cross-validation fold of the experimental transmembrane topology dataset.

| Label type | Fold 0 | Fold 1 | Fold 2 | Fold 3 | Fold 4 |
|---|---|---|---|---|---|
| β | 19 | 19 | 18 | 18 | 18 |
| Soluble | 1182 | 1182 | 1182 | 1182 | 1181 |
| α | 90 | 90 | 90 | 90 | 89 |
| α+IF | 34 | 34 | 34 | 34 | 34 |
| α+RE | 2 | 5 | 1 | 5 | 4 |
| α+RE+IF | 6 | 3 | 7 | 3 | 3 |
| α+SP | 17 | 18 | 19 | 11 | 20 |
| α+SP+IF | 2 | 1 | 0 | 6 | 0 |
| α+SP+RE | 0 | 0 | 0 | 2 | 0 |
| α+TP | 4 | 4 | 3 | 4 | 4 |
| α+TP+IF | 1 | 1 | 2 | 1 | 2 |
| SP | 184 | 184 | 184 | 183 | 183 |
| TP | 79 | 79 | 79 | 79 | 79 |

**Table S2.** Membrane type label counts per cross-validation fold of the experimental transmembrane topology dataset.

| Membrane type | Fold 0 | Fold 1 | Fold 2 | Fold 3 | Fold 4 |
|---|---|---|---|---|---|
| Archaeal membrane | 0 | 8 | 4 | 3 | 4 |
| Bacterial Gram-negative inner membrane | 39 | 38 | 39 | 45 | 17 |
| Bacterial Gram-negative outer membrane | 18 | 16 | 16 | 15 | 17 |
| Eukaryotic plasma membrane | 21 | 36 | 24 | 44 | 68 |
| Mitochondrial inner membrane | 33 | 33 | 30 | 21 | 10 |
| Endoplasmic reticulum membrane | 2 | 11 | 20 | 20 | 41 |
| Thylakoid membrane | 29 | 18 | 16 | 7 | 4 |
| Bacterial Gram-positive plasma membrane | 22 | 8 | 18 | 12 | 10 |
| Golgi membrane | 3 | 1 | 1 | 3 | 14 |
| Nuclear inner membrane | 1 | 0 | 3 | 1 | 2 |
| Endosome membrane | 1 | 2 | 1 | 2 | 7 |
| Vacuole membrane | 1 | 0 | 2 | 1 | 4 |
| Vesicle membrane | 1 | 2 | 3 | 2 | 8 |
| Viral membrane | 2 | 1 | 1 | 4 | 4 |
| Chloroplast outer membrane | 1 | 0 | 0 | 0 | 1 |
| Mitochondrial outer membrane | 1 | 3 | 4 | 3 | 2 |
| Lysosome membrane | 0 | 2 | 2 | 4 | 2 |

**Table S3.** Label counts of the UniProt membrane type multilabel dataset.

| Membrane type | Count |
|---|---|
| Archaeal membrane | 469 |
| Bacterial Gram-negative inner membrane | 832 |
| Bacterial Gram-negative outer membrane | 35 |
| Eukaryotic plasma membrane | 3585 |
| Mitochondrial inner membrane | 804 |
| Endoplasmic reticulum membrane | 3596 |
| Thylakoid membrane | 94 |
| Bacterial Gram-positive plasma membrane | 674 |
| Golgi membrane | 1779 |
| Nuclear inner membrane | 112 |
| Endosome membrane | 277 |
| Vacuole membrane | 418 |
| Vesicle membrane | 241 |
| Viral membrane | 835 |
| Chloroplast outer membrane | 41 |
| Mitochondrial outer membrane | 474 |
| Lysosome membrane | 360 |

**Table S4.** Mapping of the two modeled topological sides A and B to specific locations based on membrane type.

| Membrane type | Side A | Side B |
|---|---|---|
| Archaeal membrane | Cytoplasm | Extracellular |
| Bacterial Gram-negative inner membrane | Cytoplasm | Periplasm |
| Bacterial Gram-negative outer membrane | Extracellular | Periplasm |
| Eukaryotic plasma membrane | Cytoplasm | Extracellular |
| Mitochondrial inner membrane | Mitochondrial matrix | Intermembrane space |
| Endoplasmic reticulum membrane | Cytoplasm | Lumenal |
| Thylakoid membrane | Stroma | Thylakoid space |
| Bacterial Gram-positive plasma membrane | Cytoplasm | Periplasm |
| Golgi membrane | Cytoplasm | Lumenal |
| Nuclear inner membrane | Nucleus matrix | Intermembrane space |
| Endosome membrane | Cytoplasm | Lumenal |
| Vacuole membrane | Cytoplasm | Lumenal |
| Vesicle membrane | Cytoplasm | Lumenal |
| Viral membrane | Intravirion side | Virion surface |
| Chloroplast outer membrane | Intermembrane space | Cytoplasm |

| | | |
|---|---|---|
| Mitochondrial outer membrane | Intermembrane space | Cytoplasm |
| Lysosome membrane | Cytoplasm | Lumenal |

**Table S5.** Mapping of UniProt locations to OPM membrane types.

| UniProt location | OPM location |
|---|---|
| Chloroplast thylakoid membrane | Thylakoid membrane |
| Cellular chromatophore membrane | Thylakoid membrane |
| Virion membrane | Viral membrane |
| Mitochondrion inner membrane | Mitochondrial inner membrane |
| Basolateral cell membrane | Eukaryotic plasma membrane |
| Lateral cell membrane | Eukaryotic plasma membrane |
| Cellular thylakoid membrane | Thylakoid membrane |
| trans-Golgi network membrane | Golgi membrane |
| Nucleus inner membrane | Nuclear inner membrane |
| Endoplasmic reticulum membrane | Endoplasmic reticulum membrane |
| Clathrin-coated vesicle membrane | Vesicle membrane |
| Golgi apparatus membrane | Golgi membrane |
| Late endosome membrane | Endosome membrane |
| Flagellum membrane | Eukaryotic plasma membrane |
| Apical cell membrane | Eukaryotic plasma membrane |
| Mitochondrion | Mitochondrial inner membrane |
| Mitochondrion membrane | Mitochondrial inner membrane |
| Sarcolemma | Eukaryotic plasma membrane |
| Caveola | Eukaryotic plasma membrane |
| T-tubule | Eukaryotic plasma membrane |
| Sarcoplasmic reticulum membrane | Endoplasmic reticulum membrane |
| Lysosome membrane | Lysosome membrane |
| Ruffle membrane | Eukaryotic plasma membrane |
| Chloroplast outer membrane | Chloroplast outer membrane |
| Mitochondrion outer membrane | Mitochondrial outer membrane |
| Nucleus membrane | Nuclear inner membrane |
| Nucleus outer membrane | Nucleus outer membrane |
| Endosome | Endosome membrane |
| Postsynaptic cell membrane | Eukaryotic plasma membrane |
| Endoplasmic reticulum | Endoplasmic reticulum membrane |
| Cyanelle thylakoid membrane | Thylakoid membrane |
| Synaptic vesicle membrane | Vesicle membrane |
| Virion | Viral membrane |
| Mitochondrion matrix | Mitochondrial inner membrane |
| Cytoplasmic vesicle | Vesicle membrane |

| | |
|---|---|
| Nucleus envelope | Nuclear inner membrane |
| Rough endoplasmic reticulum membrane | Endoplasmic reticulum membrane |
| COPI-coated vesicle membrane | Vesicle membrane |
| Synaptic cell membrane | Eukaryotic plasma membrane |
| Endosome membrane | Endosome membrane |
| Postsynaptic density membrane | Eukaryotic plasma membrane |
| Cilium membrane | Eukaryotic plasma membrane |
| Microsome membrane | Endoplasmic reticulum membrane |
| Host endoplasmic reticulum membrane | Endoplasmic reticulum membrane |
| Host mitochondrion | Mitochondrial inner membrane |
| Vacuole membrane | Vacuole membrane |
| Cell outer membrane | Bacterial Gram-negative outer membrane |
| Host Golgi apparatus membrane | Golgi membrane |
| Early endosome membrane | Endosome membrane |
| Early endosome | Endosome membrane |
| Ruffle | Eukaryotic plasma membrane |
| Cytoplasmic vesicle membrane | Vesicle membrane |
| Golgi apparatus | Golgi membrane |
| Dendritic spine membrane | Eukaryotic plasma membrane |
| Host nucleus membrane | Nuclear inner membrane |
| Host rough endoplasmic reticulum membrane | Endoplasmic reticulum membrane |
| Host mitochondrion outer membrane | Mitochondrial outer membrane |
| Late endosome | Endosome membrane |
| Lysosome | Lysosome membrane |
| Host endosome | Endosome membrane |
| Host lysosome | Lysosome membrane |
| Recycling endosome membrane | Endosome membrane |
| Recycling endosome | Endosome membrane |
| Vesicle | Vesicle membrane |
| Golgi stack membrane | Golgi membrane |
| Cell tip | Eukaryotic plasma membrane |
| COPII-coated vesicle membrane | Vesicle membrane |
| Secretory vesicle membrane | Vesicle membrane |
| Multivesicular body membrane | Endosome membrane |
| Basal cell membrane | Eukaryotic plasma membrane |
| Phagosome | Vesicle membrane |
| Microvillus membrane | Eukaryotic plasma membrane |
| Chloroplast thylakoid lumen | Thylakoid membrane |
| Target cell membrane | Secreted |
| Secreted | Secreted |

**Table S6.** Metrics on the full cross-validated dataset (shown in Figure 2a).

| Type | Accuracy | F1 | Precision | Recall |
|---|---|---|---|---|
| **α** | 0.9881 ± 0.0012 | 0.94 ± 0.01 | 0.91 ± 0.01 | 0.97 ± 0.01 |
| **β** | 0.9989 ± 0.0004 | 0.95 ± 0.02 | 0.97 ± 0.02 | 0.93 ± 0.03 |
| **RE** | 0.9977 ± 0.0005 | 0.72 ± 0.06 | 0.89 ± 0.06 | 0.61 ± 0.08 |
| **IF** | 0.9876 ± 0.0012 | 0.73 ± 0.03 | 0.84 ± 0.03 | 0.64 ± 0.03 |
| **SP** | 0.9910 ± 0.0011 | 0.97 ± 0.00 | 0.99 ± 0.00 | 0.94 ± 0.01 |
| **TP** | 0.9895 ± 0.0011 | 0.90 ± 0.01 | 0.88 ± 0.02 | 0.92 ± 0.01 |

**Table S7.** Classification metrics on the TMbed cross-validated benchmark dataset (shown in Figure 2a).

| Type | Accuracy | | F1 | | Precision | | Recall | |
|---|---|---|---|---|---|---|---|---|
| | DeepTMHMM2 | TMbed | DeepTMHMM2 | TMbed | DeepTMHMM2 | TMbed | DeepTMHMM2 | TMbed |
| **α** | 0.9930 ± 0.0015 | **0.9960 ± 0.0011** | 0.96 ± 0.01 | **0.98 ± 0.01** | 0.94 ± 0.01 | **0.99 ± 0.01** | **0.98 ± 0.01** | 0.97 ± 0.01 |
| **β** | **0.9991 ± 0.0005** | 0.9988 ± 0.0006 | **0.97 ± 0.02** | 0.96 ± 0.02 | 1.00 ± 0.00 | 1.00 ± 0.00 | **0.94 ± 0.03** | 0.93 ± 0.04 |
| **SP** | 0.9921 ± 0.0015 | **0.9970 ± 0.0010** | 0.96 ± 0.01 | **0.99 ± 0.00** | **1.00 ± 0.00** | 0.99 ± 0.01 | 0.93 ± 0.01 | **0.99 ± 0.01** |

**Table S8.** Classification metrics on the DeepTMHMM 1 cross-validated benchmark dataset (shown in Figure 2a). (v2 = DeepTMHMM2, v1 = DeepTMHMM 1)

| Type | Accuracy | | F1 | | Precision | | Recall | |
|---|---|---|---|---|---|---|---|---|
| | v2 | v1 | v2 | v1 | v2 | v1 | v2 | v1 |
| **α** | **0.9899 ± 0.0027** | 0.9884 ± 0.0029 | 0.95 ± 0.01 | 0.95 ± 0.01 | **0.92 ± 0.02** | 0.91 ± 0.02 | **0.99 ± 0.01** | 0.98 ± 0.01 |
| **β** | **0.9993 ± 0.0007** | 0.9986 ± 0.0010 | **0.99 ± 0.01** | 0.98 ± 0.01 | **1.00 ± 0.00** | 0.98 ± 0.02 | 0.98 ± 0.02 | 0.98 ± 0.02 |
| **SP** | 0.9754 ± 0.0042 | **0.9899 ± 0.0027** | 0.96 ± 0.01 | **0.98 ± 0.00** | **1.00 ± 0.00** | 0.99 ± 0.00 | 0.91 ± 0.01 | **0.97 ± 0.01** |

**Table S9.** Classification metrics on the SignalP cross-validated benchmark dataset (shown in Figure 2a).

| Type | Accuracy | | F1 | | Precision | | Recall | |
|---|---|---|---|---|---|---|---|---|
| | DeepTMHMM2 | SignalP 6.0 | DeepTMHMM2 | SignalP 6.0 | DeepTMHMM2 | SignalP 6.0 | DeepTMHMM2 | SignalP 6.0 |
| **SP** | 0.9900 ± 0.0014 | **0.9925 ± 0.0012** | 0.96 ± 0.01 | **0.97 ± 0.00** | **1.00 ± 0.00** | 0.99 ± 0.00 | 0.92 ± 0.01 | **0.95 ± 0.01** |

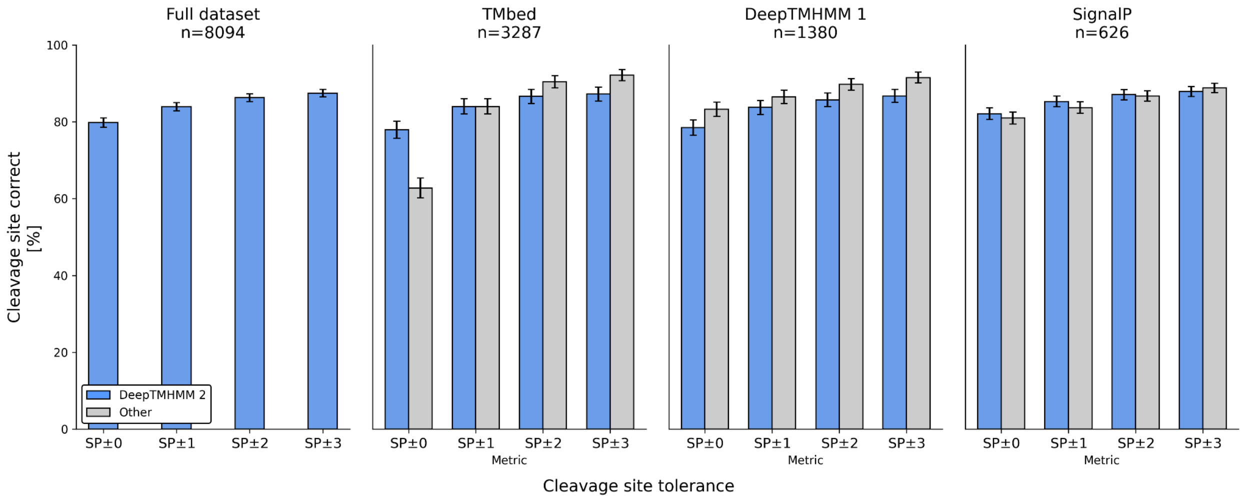


**Figure S2.** % correct for SP prediction at varying tolerance windows around the true cleavage site.

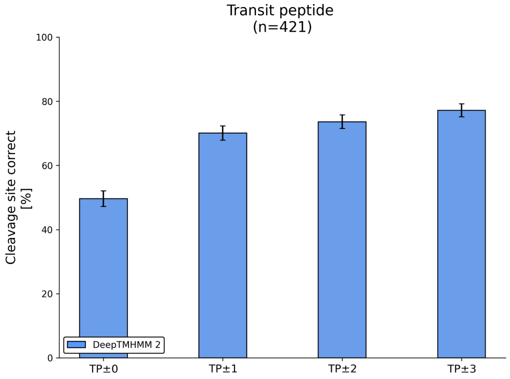


**Figure S3.** % correct for TP prediction at varying tolerance windows around the true cleavage site.

**Table S10.** The full list of methods (in alphabetical order) that were included in the benchmark of DeepTMHMM2. Sequence representation denotes if a method only operates on a single amino acid sequence or leverages evolutionary information via profiles from multiple sequence alignment or embeddings. Consensus methods ensemble multiple prediction models. Methods that do not perform complete topology prediction or whose web server is unavailable were excluded (TMPSS[13], HDNNtopss[14], MASSP[15]).

| Method | Server URL/Repository | Year | Protein type | Sequence representation | SP prediction | Consensus method |
|---|---|---|---|---|---|---|
| BetAware-Deep[16] | https://busca.biocomp.unibo.it/betaware2/ | 2021 | beta barrels | Evolutionary information | N | N |
| BOCTOPUS2[17] | Htt ps://b2.topcons.net | 2016 | beta barrels | Evolutionary information | N | N |
| CCTOP[18] | http://cctop.ttk.hu | 2015 | alpha helical | Evolutionary information | Y | Y |
| DeepTMpred[19] | https://github.com/ISYSLAB-HUST/DeepTMpred/ | 2022 | alpha helical | Evolutionary information | Y | Y |
| HMM-TM[20] | http://bioinformatics.biol.uoa.gr/HMM- TM/ | 2006 | alpha helical | Single sequence | N | N |
| HMMpTM[21] | http://biophysics.biol.uoa.gr/HMMpTM/ | 2013 | alpha helical | Single sequence | N | N |
| HMMTOP[22] | http://www.enzim.hu/hmmtop/ | 2001 | alpha helical | Single sequence | N | N |
| MemBrain 3.0/3.1[23,24] | http://www.csbio.sjtu.edu.cn/bioinf/MemBrain/ | 2020 | alpha helical | Evolutionary information | N | N |
| MEMSAT3[25] | http://bioinfadmin.cs.ucl.ac.uk/downloads/ memsat/ | 2007 | alpha helical | Evolutionary information | N | N |
| MEMSAT-SVM[26] | http://bioinfadmin.cs.ucl.ac.uk/downloads/ memsat-svm/ | 2009 | alpha helical | Evolutionary information | Y | N |
| OCTOPUS[27] | http://octopus.cbr.su.se/ | 2008 | alpha helical | Evolutionary information | N | N |
| Philius[28] | http://www.yeastrc.org/philius/pages/philiu s/runPhilius.jsp | 2008 | alpha helical | Single sequence | Y | N |

| Method | Server URL/Repository | Year | Protein type | Sequence representation | SP prediction | Consensus method |
|---|---|---|---|---|---|---|
| Phobius[29] | http://phobius.sbc.su.se/ | 2004 | alpha helical | Single sequence | Y | N |
| PolyPhobius[30] | http://phobius.sbc.su.se/ | 2005 | alpha helical | Evolutionary information | Y | N |
| PRED-TMBB2[31] | http://www.compgen.org/tools/PRED- TMBB2 | 2016 | beta barrels | Evolutionary information | Y | N |
| PRED-TMSdeep[32] | https://hannibal.dib.uth.gr/PRED-TMSdeep/ | 2026 | both | Evolutionary information | Y | N |
| SCAMPI2[33] | http://scampi.bioinfo.se/ | 2016 | alpha helical | Both | N | N |
| SPOCTOPUS[34] | http://octopus.cbr.su.se/ | 2008 | alpha helical | Evolutionary information | Y | N |
| TMbed[35] | https://github.com/BernhoferM/TMbed | 2022 | both | Evolutionary information | Y | N |
| TMHMM 2.0[36] | https://services.healthtech.dtu.dk/services/TMHMM-2.0/ | 2001 | alpha helical | Single sequence | N | N |
| TOPCONS[37] | http://topcons.net/ | 2015 | alpha helical | Evolutionary information | Y | Y |

**Table S11.** Extended benchmark for TM-α prediction performance.

| Method | F1 α-TM | F1 SP | % correct (incl. SP) | % correct (excl. SP) | % correct SP±0 | % correct SP±1 | % correct SP±2 | % correct SP±3 |
|---|---|---|---|---|---|---|---|---|
| DeepTMHMM2 | **0.94** | **0.97** | **85.6** | **87.2** | **79.8** | **83.9** | **86.3** | **87.5** |
| CCTOP | 0.83 | 0.18 | 85.3 | 86.2 | 5.8 | 8.0 | 8.6 | 9.0 |
| DeepTMpred* | 0.40 | - | - | 42.7 | - | - | - | - |
| HMM-TM | 0.28 | - | - | 38.1 | - | - | - | - |
| HMMpTM | 0.31 | - | - | 48.7 | - | - | - | - |
| MemBrain 3.0 | 0.37 | 0.98** | 61.8 | 63.2 | 85.6** | 89.2** | 91.7** | 92.9** |
| MEMSAT3 | 0.62 | - | - | 69.2 | - | - | - | - |
| MEMSAT-SVM | 0.56 | 0.42 | 55.0 | 63.8 | 2.4 | 3.6 | 5.9 | 8.3 |
| OCTOPUS | 0.65 | - | - | 56.7 | - | - | - | - |
| Philius | 0.85 | 0.82 | 58.7 | 60.8 | 53.5 | 62.1 | 69.9 | 78.0 |
| Phobius | 0.82 | 0.87 | 53.8 | 55.6 | 52.7 | 61.9 | 69.0 | 77.1 |
| PolyPhobius | 0.78 | 0.89 | 59.0 | 61.0 | 43.6 | 55.4 | 66.5 | 78.1 |
| SCAMPI2 MSA | 0.76 | - | - | 57.3 | - | - | - | - |
| SCAMPI2 single | 0.72 | - | - | 47.2 | - | - | - | - |
| SPOCTOPUS | 0.56 | 0.84 | 55.4 | 57.3 | 35.3 | 48.5 | 60.3 | 70.4 |
| TMHMM | 0.83 | - | - | 50.6 | - | - | - | - |
| TOPCONS2 | 0.83 | 0.89 | 64.7 | 67.8 | 27.2 | 39.0 | 51.2 | 64.3 |
| AF2+TMDET 4.0*** | 0.65 | - | - | 83.3 | - | - | - | - |
| AF2+ PPM 2.0*** | 0.90 | - | - | 68.3 | - | - | - | - |

*The DeepTMpred predictor reports sides *1* and *2* without defining a correspondence to the standard inside-outside terminology used in the DeepTMpred paper. We map as *1*=I and *2*=O, which gives a higher performance than the inverse.
**MemBrain 3.0 was run with Signal-3L 4.0 for SP prediction, which was trained on the SignalP 6.0-derived data used for testing here.
***The % correct metric excludes the N-terminal a/b-I/O orientation criterion for structure membrane insertion algorithms.

**Table S12.** Extended benchmark for TM-β prediction performance.

| Method | F1 β-TM | % correct (incl. SP) | % correct (excl. SP) |
|---|---|---|---|
| DeepTMHMM2 | **0.54** | **76.1** | **83.7** |
| BetAware-Deep | 0.32 | - | 64.1 |
| BOCTOPUS2 | 0.29 | - | 66.3 |
| PRED-TMBB2 | 0.25 | 47.8 | 54.3 |
| AF2+TMDET 4.0** | 0.84 | - | 64.1 |
| AF2+PPM 2.0** | 0.90 | - | 32.6 |

**The % correct metric excludes the N-terminal a/b-I/O orientation criterion for structure membrane insertion algorithms.

**Table S13.** Extended benchmark for RE prediction performance. For MemBrain 3.0, RE refer to what is named *half-TMH* in the original paper.

| Method | Type | | | Topology |
|---|---|---|---|---|
| | F1 | Precision | Recall | % correct |
| DeepTMHMM2 | **0.72** | **0.89** | 0.61 | **58.5** |
| MemBrain 3.0 | 0.01 | 0.01 | **0.85** | 9.8 |
| MEMSAT-SVM | 0.38 | 0.83 | 0.24 | 19.5 |
| AF2 + TMDET 4.0 | 0.39 | 0.27 | **0.73** | 51.2 |

**Table S14.** Extended benchmark for IF prediction performance. To match MemBrain 3.1, IF predictions in soluble proteins are ignored when evaluating false positives in this analysis.

| Method | Type | | | Topology |
|---|---|---|---|---|
| | F1 | Precision | Recall | % correct |
| DeepTMHMM2 | **0.74** | **0.88** | 0.64 | **43.3** |
| MemBrain 3.1 | 0.31 | 0.19 | **0.86** | 28.8 |
| AF2 + TMDET 4.0 | 0.39 | 0.82 | 0.25 | 15.3 |

**Table S15.** Extended benchmark for PRED-TMSdeep, using cross-validated prediction common sets.

| Type | n | F1 | | % correct | |
|---|---|---|---|---|---|
| | | DeepTMHMM2 | PRED-TMSdeep | DeepTMHMM2 | PRED-TMSdeep |
| **α** | 452 | 0.97 ± 0.01 | **0.99 ± 0.00** | **87.8 ± 1.5** | 85.8 ± 1.6 |
| **β** | 74 | **0.97 ± 0.02** | 0.90 ± 0.03 | **79.7 ± 4.7** | 71.6 ± 5.2 |
| **SP** | 533 | 0.96 ± 0.01 | **0.99 ± 0.00** | **80.5 ± 1.7** | 64.4 ± 2.1 |

**Table S16.** Membrane type prediction performance.

| Membrane type | MCC | AUPRC | AUROC | F1 | % positive | # positive |
|---|---|---|---|---|---|---|
| Archaeal membrane | 0.766 | 0.808 | 0.977 | 0.765 | 2.2% | 19 |
| Bacterial Gram-negative inner membrane | 0.709 | 0.857 | 0.955 | 0.768 | 20.4% | 178 |
| Bacterial Gram-negative outer membrane | 0.939 | 0.980 | 0.996 | 0.944 | 9.4% | 82 |
| Eukaryotic plasma membrane | 0.857 | 0.941 | 0.983 | 0.889 | 22.1% | 193 |
| Mitochondrial inner membrane | 0.882 | 0.943 | 0.973 | 0.897 | 14.6% | 127 |
| Endoplasmic reticulum membrane | 0.558 | 0.643 | 0.904 | 0.607 | 10.8% | 94 |
| Thylakoid membrane | 0.633 | 0.796 | 0.962 | 0.645 | 8.5% | 74 |
| Bacterial Gram-positive plasma membrane | 0.575 | 0.577 | 0.938 | 0.606 | 8.0% | 70 |
| Golgi membrane | 0.240 | 0.188 | 0.877 | 0.160 | 2.5% | 22 |
| Nuclear inner membrane | 0.142 | 0.068 | 0.832 | 0.129 | 0.8% | 7 |
| Endosome membrane | 0.195 | 0.074 | 0.878 | 0.132 | 1.5% | 13 |
| Vacuole membrane | 0.256 | 0.136 | 0.886 | 0.250 | 0.9% | 8 |
| Vesicle membrane | 0.336 | 0.244 | 0.901 | 0.345 | 1.8% | 16 |
| Viral membrane | 0.505 | 0.463 | 0.935 | 0.500 | 1.4% | 12 |
| Chloroplast outer membrane | 0.407 | 0.667 | 0.998 | 0.400 | 0.2% | 2 |
| Mitochondrial outer membrane | 0.448 | 0.359 | 0.766 | 0.421 | 1.5% | 13 |
| Lysosome membrane | 0.167 | 0.068 | 0.897 | 0.090 | 1.2% | 10 |

### Bibliography


1. Tusnády, G. E., Dosztányi, Z. & Simon, I. TMDET: web server for detecting transmembrane regions of proteins by using their 3D coordinates. *Bioinformatics* **21**, 1276–1277 (2005).
2. Kozma, D., Simon, I. & Tusnády, G. E. PDBTM: Protein Data Bank of transmembrane proteins after 8 years. *Nucleic Acids Res.* **41**, D524-529 (2013).
3. Tusnády, G. E. & Gerdán, C. TmDet 4.0: determining membrane orientation of transmembrane proteins from 3D structure. *Nucleic Acids Res.* **53**, W542–W546 (2025).
4. Erdogmus, S. *et al.* Helix 8 is the essential structural motif of mechanosensitive GPCRs. *Nat. Commun.* **10**, 5784 (2019).
5. Dall'Osto, L., Bressan, M. & Bassi, R. Biogenesis of light harvesting proteins. *Biochim. Biophys. Acta* **1847**, 861–871 (2015).
6. Steinegger, M. & Söding, J. MMseqs2 enables sensitive protein sequence searching for the analysis of massive data sets. *Nat. Biotechnol.* **35**, 1026–1028 (2017).
7. van Kempen, M. *et al.* Fast and accurate protein structure search with Foldseek. *Nat. Biotechnol.* **42**, 243–246 (2024).
8. Varadi, M. *et al.* AlphaFold Protein Structure Database in 2024: providing structure coverage for over 214 million protein sequences. *Nucleic Acids Res.* **52**, D368–D375 (2023).
9. Lin, Z. *et al.* Evolutionary-scale prediction of atomic-level protein structure with a language model. *Science* **379**, 1123–1130 (2023).
10. Zhang, Y. & Skolnick, J. Scoring function for automated assessment of protein structure template quality. *Proteins* **57**, 702–710 (2004).
11. Lomize, M. A., Pogozheva, I. D., Joo, H., Mosberg, H. I. & Lomize, A. L. OPM database and PPM web server: resources for positioning of proteins in membranes. *Nucleic Acids Res.* **40**, D370-376 (2012).
12. Dobson, L. *et al.* TmAlphaFold database: membrane localization and evaluation of AlphaFold2 predicted alpha-helical transmembrane protein structures. *Nucleic Acids Res.* **51**, D517–D522 (2023).
13. Liu, Z. *et al.* TMPSS: A Deep Learning-Based Predictor for Secondary Structure and Topology Structure Prediction of Alpha-Helical Transmembrane Proteins. *Front. Bioeng. Biotechnol.* **8**, 629937 (2020).
14. Gao, T., Zhao, Y., Zhang, L. & Wang, H. Secondary and Topological Structural Merge Prediction of Alpha-Helical Transmembrane Proteins Using a Hybrid Model Based on Hidden Markov and Long Short-Term Memory Neural Networks. *Int. J. Mol. Sci.* **24**, 5720 (2023).
15. Li, B., Mendenhall, J., Capra, J. A. & Meiler, J. A Multitask Deep-Learning Method for Predicting Membrane Associations and Secondary Structures of Proteins. *J. Proteome Res.* **20**, 4089–4100 (2021).
16. Madeo, G., Savojardo, C., Martelli, P. L. & Casadio, R. BetAware-Deep: An Accurate Web Server for Discrimination and Topology Prediction of Prokaryotic Transmembrane β-barrel Proteins. *J. Mol. Biol.* **433**, 166729 (2021).
17. Hayat, S., Peters, C., Shu, N., Tsirigos, K. D. & Elofsson, A. Inclusion of dyad-repeat pattern improves topology prediction of transmembrane β-barrel proteins. *Bioinformatics* **32**, 1571–1573 (2016).
18. Dobson, L., Reményi, I. & Tusnády, G. E. CCTOP: a Consensus Constrained TOPology prediction web server. *Nucleic Acids Res.* **43**, W408-412 (2015).
19. Wang, L., Zhong, H., Xue, Z. & Wang, Y. Improving the topology prediction of α-helical transmembrane proteins with deep transfer learning. *Comput. Struct. Biotechnol. J.* **20**, 1993–2000 (2022).
20. Bagos, P. G., Liakopoulos, T. D. & Hamodrakas, S. J. Algorithms for incorporating prior topological information in HMMs: application to transmembrane proteins. *BMC Bioinformatics* **7**, 189 (2006).

21. Tsaousis, G. N., Bagos, P. G. & Hamodrakas, S. J. HMMpTM: improving transmembrane protein topology prediction using phosphorylation and glycosylation site prediction. *Biochim. Biophys. Acta* **1844**, 316–322 (2014).
22. Tusnády, G. E. & Simon, I. The HMMTOP transmembrane topology prediction server. *Bioinformatics* **17**, 849–850 (2001).
23. Feng, S.-H., Zhang, W.-X., Yang, J., Yang, Y. & Shen, H.-B. Topology Prediction Improvement of α-helical Transmembrane Proteins Through Helix-tail Modeling and Multiscale Deep Learning Fusion. *J. Mol. Biol.* **432**, 1279–1296 (2020).
24. Feng, S.-H., Xia, C.-Q., Zhang, P.-D. & Shen, H.-B. Ab-Initio Membrane Protein Amphipathic Helix Structure Prediction Using Deep Neural Networks. *IEEE/ACM Trans. Comput. Biol. Bioinform.* **19**, 795–805 (2022).
25. Jones, D. T. Improving the accuracy of transmembrane protein topology prediction using evolutionary information. *Bioinformatics* **23**, 538–544 (2007).
26. Nugent, T. & Jones, D. T. Transmembrane protein topology prediction using support vector machines. *BMC Bioinformatics* **10**, 159 (2009).
27. Viklund, H. & Elofsson, A. OCTOPUS: improving topology prediction by two-track ANN-based preference scores and an extended topological grammar. *Bioinformatics* **24**, 1662–1668 (2008).
28. Reynolds, S. M., Käll, L., Riffle, M. E., Bilmes, J. A. & Noble, W. S. Transmembrane Topology and Signal Peptide Prediction Using Dynamic Bayesian Networks. *PLOS Comput. Biol.* **4**, e1000213 (2008).
29. Käll, L., Krogh, A. & Sonnhammer, E. L. L. A Combined Transmembrane Topology and Signal Peptide Prediction Method. *J. Mol. Biol.* **338**, 1027–1036 (2004).
30. Käll, L., Krogh, A. & Sonnhammer, E. L. L. An HMM posterior decoder for sequence feature prediction that includes homology information. *Bioinformatics* **21**, i251–i257 (2005).
31. Tsirigos, K. D., Elofsson, A. & Bagos, P. G. PRED-TMBB2: improved topology prediction and detection of beta-barrel outer membrane proteins. *Bioinformatics* **32**, i665–i671 (2016).
32. Moschos, G. A., Tsirigos, K. D., Tamposis, I. A. & Bagos, P. G. PRED-TMSdeep: Prediction of Transmembrane Topology and Signal Peptides Using Deep Learning. *Biology* **15**, 1016 (2026).
33. Peters, C., Tsirigos, K. D., Shu, N. & Elofsson, A. Improved topology prediction using the terminal hydrophobic helices rule. *Bioinformatics* **32**, 1158–1162 (2016).
34. Viklund, H., Bernsel, A., Skwark, M. & Elofsson, A. SPOCTOPUS: a combined predictor of signal peptides and membrane protein topology. *Bioinformatics* **24**, 2928–2929 (2008).
35. Bernhofer, M. & Rost, B. TMbed: transmembrane proteins predicted through language model embeddings. *BMC Bioinformatics* **23**, 326 (2022).
36. Krogh, A., Larsson, B., von Heijne, G. & Sonnhammer, E. L. Predicting transmembrane protein topology with a hidden Markov model: application to complete genomes. *J. Mol. Biol.* **305**, 567–580 (2001).
37. Tsirigos, K. D., Peters, C., Shu, N., Käll, L. & Elofsson, A. The TOPCONS web server for consensus prediction of membrane protein topology and signal peptides. *Nucleic Acids Res.* **43**, W401-407 (2015).
